# *In extracto* cryo-EM reveals eEF2 as a major hibernation factor on 60S and 80S particles

**DOI:** 10.1101/2025.11.25.690450

**Authors:** Zahra Seraj, Ximena Zottig, Chun-Ying Huang, Anna B. Loveland, Stephen Diggs, Emily Sholi, Nikolaus Grigorieff, Andrei A. Korostelev

## Abstract

Cryogenic electron microscopy (cryo-EM) made impressive progress in resolving cellular macromolecules and their detailed interactions. Single-particle cryo-EM traditionally relies on purified macromolecules and lacks the complexity of cellular environments, whereas *in situ* cryo-EM or cryo-ET require extensive sample preparation and data acquisition, presenting challenges in achieving high resolution. We describe cryo-EM of cellular lysates—*in extracto* cryo-EM—allowing the flexibility and high-resolution of cryo-EM in the context of cellular components. High-resolution 2D template matching (2DTM) yields ∼2.2 Å maps of the mammalian translational apparatus. Elongating ribosome abundances in primate cell lines (MCF-7 and BSC-1) and rabbit reticulocyte lysates range from ∼70% to ∼10%, reflecting translational stress responses. Non-translating (hibernating) ribosomes carrying no mRNA, feature numerous proteins shielding ribosomal functional centers. Elongation factor 2 (eEF2) is the most abundant hibernation factor bound to >95% of ribosomes and, unexpectedly, to 60S subunits. eEF2•GDP is stabilized by interactions with the sarcin-ricin loop and protein uL14. Hibernating ribosomes also feature LARP1 involved in initiation and mTOR signaling; eIF5A implicated in elongation and termination; and other factors, exposing the variety of hibernation scenarios. Our work underscores the efficiency and potential of *in extracto* cryo-EM to discover native cellular complexes and mechanisms at near-atomic resolution.

## Introduction

The renaissance of structural biology, thanks to high-resolution cryogenic electron microscopy (cryo-EM) [1–3], is rapidly expanding our understanding of biological processes, from visualizing dynamic macromolecules to discovering new cellular compartments [4–7]. Transmission electron microscopy is employed to study biological samples in two major applications: single-particle (*in vitro*) cryo-EM of purified macromolecules or reconstituted complexes, and cellular/tissue (*in situ*) cryo-EM, the most notable approach being cryogenic electron tomography (cryo-ET). Each approach has advantages and limitations. Single-particle cryo-EM allows elucidation of structural details of interactions and reactions. Here, analyses of large datasets (hundreds of thousands to >1 million of particles) separate conformationally and compositionally distinct states (“classes”), yielding near-atomic-resolution (∼2 to 3.5 Å) insights into macromolecular dynamics and interactions. Furthermore, datasets can be collected at different time points of a reaction, allowing time-resolved cryo-EM to uncover transient intermediates in the absence of inhibitors [8–10]. For example, recent studies revealed the dynamics of the ribosome and translation factors during translation initiation [11], elongation [12, 13], termination [14, 15], and recycling [16, 17]. However, because samples are usually assembled from a limited set of purified components, this approach does not account for yet-to-be-discovered cellular interactions that may be important for the processes under investigation. To fill that gap, electron microscopy can be performed on cellular samples. Cryo-ET is one of the most popular approaches, providing impressive three-dimensional views of cellular compartments and large macromolecular complexes *in situ* [18] [19] [20]. Because the cellular environment is densely filled with macromolecules, electron microscopy of cellular samples remains a challenge and requires laborious sample preparation, data collection and processing [18]. For example, focused ion beam (FIB) milling of frozen cells to generate the number of thin areas (lamellae) necessary to collect large datasets can take days if not weeks. Furthermore, sub-tomogram averaging of specific macromolecular densities usually does not achieve the near-atomic resolution of similar macromolecules analyzed by single-particle cryo-EM. Finally, it remains challenging to manipulate cells to study a particular aspect of a cellular process, such as a specific step of a multi-step pathway, without using modulators/inhibitors that may bias cells off pathway.

In this work, we tested whether the advantages of both methods can be combined by exploiting cellular extracts to elucidate novel cellular complexes and interactions at high resolution. We demonstrate that high-resolution 2D template matching (2DTM) [21] enables the reconstructions of mammalian 80S complexes at ∼2.2 Å resolution, similar to single-particle cryo-EM from purified ribosomes. We studied 80S translation complexes in several mammalian lysates, starting with the optimization of the cryo-EM/2DTM pipeline on lysates obtained from a human breast cancer cell line MCF-7 and a monkey kidney cell line BSC-1. Our aim was to compare the translational states of ribosomes in distinct cell types and to visualize how cellular stress affects these states in MCF-7. We find that MCF-7 and BSC-1 lysates feature 60-70% of elongation-like complexes, echoing the analyses of mammalian translation in cells [22, 23]. In nutrient-deprived MCF-7, the abundance of elongation complexes is reduced by ∼15%, consistent with inhibition of translation upon eIF2α phosphorylation [24, 25].

We also characterized the widely used rabbit reticulocyte lysates (RRL), the predominant model system to study translation since the 1970s [26]. RRL is a flexible and tunable system that allows monitoring of translation of a single mRNA (e.g, a reporter mRNA encoding a luciferase). Studies in RRL have led to discoveries or mechanistic descriptions of translation factors [27], internal ribosome entry sites [28, 29], and other translational regulators [30, 31]. Yet, the composition of translation complexes in RRL remains uncharacterized. Unlike purified ribosome complexes, we find more multi-part compositions of 80S complexes, bringing new insights into translation regulation. For example, we find that after a few minutes of translation in RRL, only a small fraction of ribosomes is involved in translating a reporter mRNA (10-12%), whereas the majority of 80S complexes are in “hibernating” complexes bound with eEF2 but lacking mRNA. In addition, these complexes contain eIF5A, primarily implicated in translation elongation and termination, La-related protein 1 (LARP1) implicated in regulation of terminal oligopyrimidine (TOP)-containing mRNAs [32], and previously identified hibernation factors SERBP1, CCDC124 and IFRD2 [33–35]. These findings reveal that all functional centers of the ribosome are shielded in 80S hibernating complexes.

Simple lysate preparation and straightforward data processing make “*in extracto* cryo-EM*”* an efficient approach, allowing visualization of cellular complexes at near-atomic resolution. Furthermore, some parameters, such as substrate (*e.g.,* mRNA) composition and concentration, can be tuned to assemble the desired complex(es) without the need to reconstitute the specific complex(es) or to perturb cells by modulators/inhibitors. Our approach therefore offers a flexible method to study translation, and potentially other processes, in the context of authentic cellular components that may affect these processes.

## Results and Discussion

### 2DTM resolves translating and hibernating ribosomes in fresh cell lysates

In our recent work describing angiogenin activation by ribosomes in rabbit reticulocyte lysates (RRL) [36], we investigated how the angiogenin-ribosome interaction is affected by other cellular components. Typical lysate preparations produce dense and viscous samples, making it challenging to prepare cryo-EM grids with thin sample layers and to analyze them by standard cryo-EM methods. Indeed, standard data-processing workflows using Relion and *cis*TEM [37, 38], which we and others routinely use to process single-particle data, failed to yield high-quality maps of distinct classes. Extensive culling of micrographs, manual particle picking, and 2D classification to remove low-quality particles was required, but this eventually resulted in small, good-quality particles stacks (∼6 or ∼17 thousand) yielding only three distinct ribosome classes at ∼3.2-4.4 Å resolution. To overcome this problem, we used high-resolution two-dimensional template matching (2DTM), which recently was shown to successfully identify ribosomes in dense cellular environments [21]. This method substantially improved the number of picked particles (∼84,000), yielding higher-resolution maps (3 Å), and separating tens of classes from the same dataset (Methods and Table S1).

Inspired by the success of the 2DTM approach on a previously published dataset [36], we used this approach on freshly prepared mammalian cell lysates. We analyzed the breast-cancer-derived MCF-7 cell line under normal and nutrient-deprived conditions, testing whether the expected downregulation of the translational apparatus can be observed in lysates. We also analyzed the African Green Monkey kidney cell line BSC-1 under normal growth conditions to compare the distribution of translational complexes in distinct organisms and tissues.

We use a mild lysis procedure (Fig. 1A) relying on permeabilization of cellular membranes by the non-ionic detergent digitonin [39–41], allowing preservation of translation-competent ribosome states [42]. In the first approach, MCF-7 cells were permeabilized by a digitonin-containing lysis buffer, and the cytosol was collected from cell-containing wells and applied to cryo-EM grids. In the second approach, BSC-1 cells were detached from wells, incubated in the lysis buffer, briefly centrifuged to remove cell debris, and the supernatant was applied to cryo-EM grids. Preparation of the lysate in both cases was fast (under 10 minutes) and the resulting grids were directly suitable for cryo-EM data collection. In cryo-EM images, collected using standard data collection procedures employed in single-particle EM, cellular filaments can be seen next to ribosomes in the cytosol of permeabilized cells (**Figure 1—figure supplement 1**). Even in the lysates separated from cell debris, additional cellular components are readily visible, including mitochondria (**Figure 1—figure supplement 1**). Thus, a simple and streamlined approach can be used to visualize ribosomes and other cellular components.

**Figure 1.**
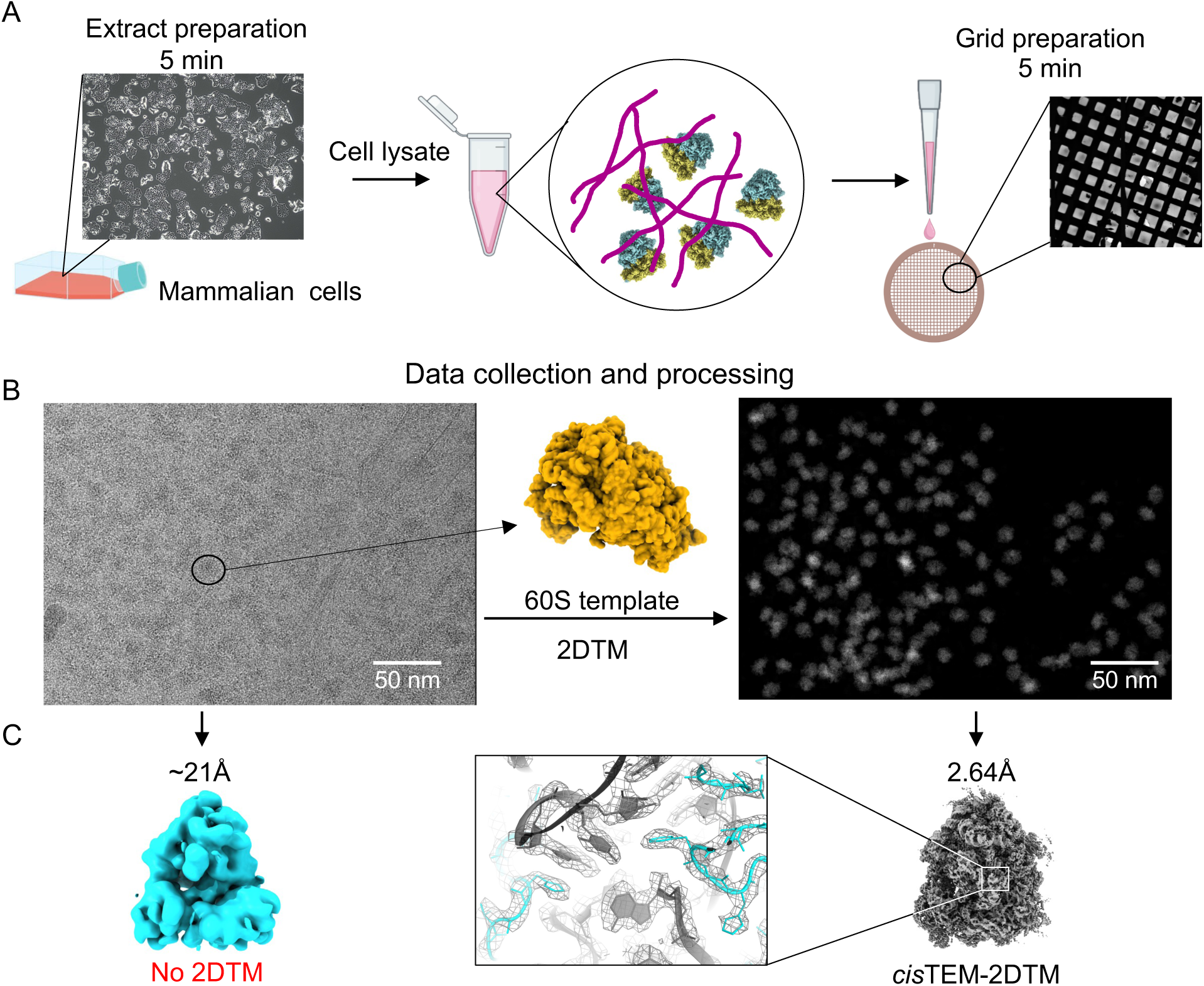
*In extracto* cryo-EM workflow for sample preparation and data collection. **A**) Sample and grid preparation from primate semi-permeabilized cells. Examples of an MCF-7 cell culture (left) and grid with cell lysate (right) are shown. **B**) Representative lysate micrograph (left) and 2DTM processing yielding positions and orientations of 60S subunits (right). **C**) Examples of averaged maps (prior to classification) from typical single-particle picking pipelines (cyan) and from 2DTM processing (middle and gray). The close-up view highlights densities of 28S rRNA nucleotides (gray) and ribosomal protein residues (cyan). Also see **Figure 2—figure supplement 1**.

Particles were picked in the collected datasets using a high-resolution template obtained from a vacant (i.e. without translation factors) mammalian 60S ribosomal subunit [43], and they were classified, with 3D maximum-likelihood classification and a focus mask on the ribosomal A site (Methods; **Figure 2—figure supplement 1-3**). Classification yielded primarily 80S ribosomes, and most 60S particles contained additional bound factors, underscoring that picking with the vacant 60S-template yielded largely template-unbiased results (**Figure 2—figure supplement 1**). This result echoes our concurrent finding that using 40S or partial 40S templates yields a variety of initiation complexes and 80S classes, revealing densities beyond those in the template [44]. The reconstruction of individual maps achieved the resolution of ∼2.2 Å for the larger BSC-1 dataset (∼525,000 particles) and ∼3 Å for each MCF-7 dataset featuring a much lower number of particles (∼80,000).

Data obtained from MCF-7 and BSC-1 grown under normal growth conditions, yielded predominantly translation elongation states containing mRNA and tRNAs (∼70% in MCF-7 and ∼60% in BSC-1, Table S2). Most of these particles are in an mRNA-decoding state, containing canonical eukaryotic elongation factor 1A (eEF1A) with A/T tRNA (A/Ternary complex) docked into the small-subunit A site or with density resembling extended-eEF1A *[22]* with A, P and E tRNAs (Fig. 2A, B, R). The remaining elongation classes contain mRNA with at least two tRNAs within a non-rotated or rotated (post-translocation) ribosome. A translocated 80S ribosome contains eEF2 in the A site, in the presence of P and E tRNAs, similar to the post-translocation state (POST-3: PDB ID 6GZ5) *[45]* (Fig. 2E). These maps closely resemble the structures of cellular elongation-state ribosomes identified in other cell types via cryo-ET *[22, 46, 47]* or cryo-EM *[12, 42, 48, 49]*. Notably, all ribosome and 60S classes contain ErbB3-binding protein 1 (EBP1) bound next to the polypeptide exit tunnel (**Figure 3—figure supplement 1**). Previously implicated in transiently binding hibernating/vacant ribosomes and regulating translation under stress *[50–52]*, EBP1 appears to be a resident protein on both translating and non-translating ribosome populations.

**Figure 2.**
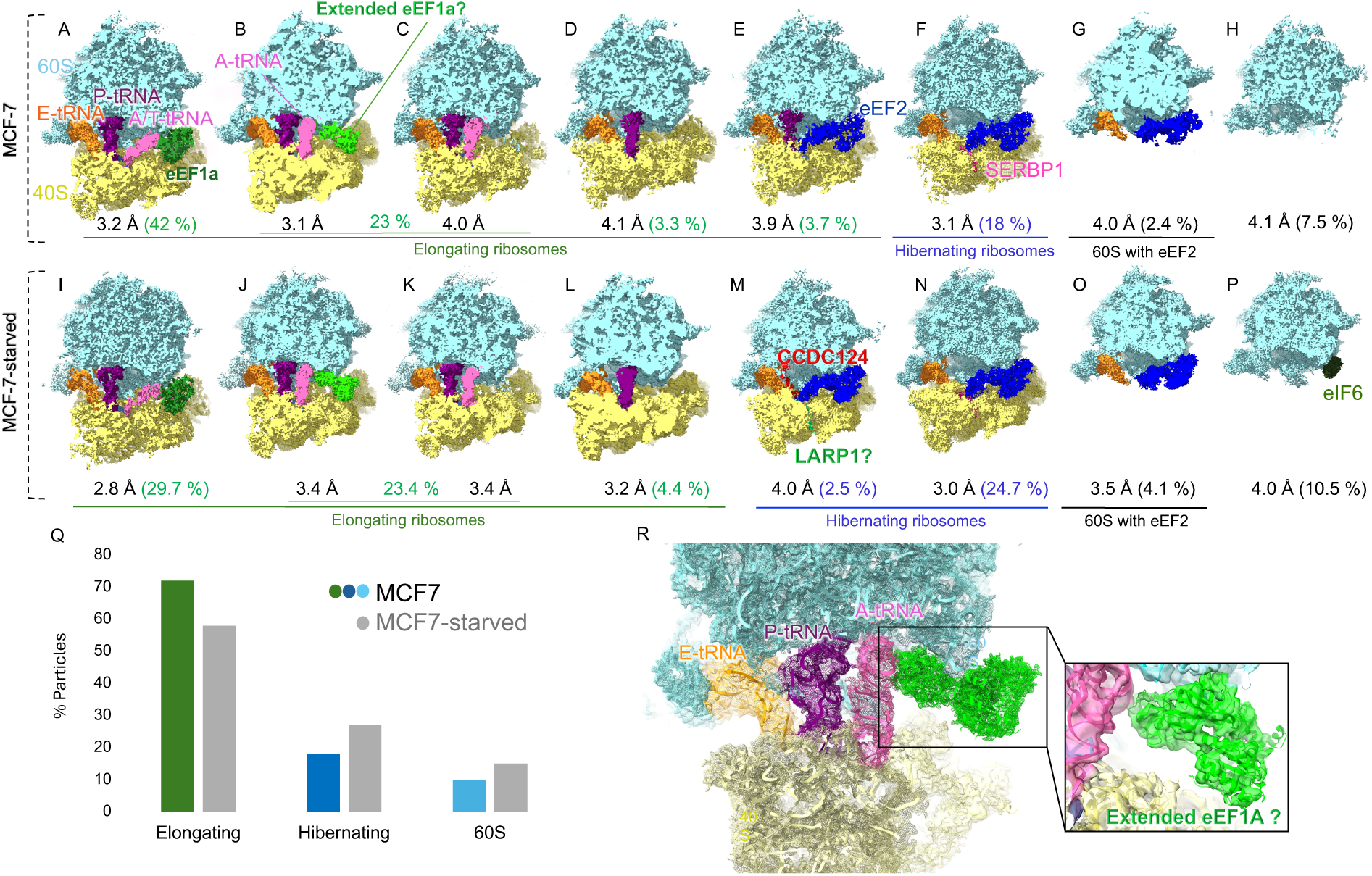
Ribosome and 60S particle distributions in control MCF-7 and starved MCF-7 cells. **A-H**) Eight cryo-EM maps correspond to elongating ribosomes (A-E): codon sampling with eEF1A and A/T tRNA (A), non-rotated with A/A, P/P,E/E tRNAs and putative extended eEF1A (B), non-rotated pre-translocation with A/A, P/P,E/E tRNAs (C), rotated pre-translocation with hybrid-state A/P and P/E tRNAs (D), and post-translocation ribosome with eEF2, P and E tRNAs (E); hibernating rotated ribosome with eEF2, SERBP1 and P/E tRNA (F); 60S subunits with eEF2 and E-tRNA (G); and vacant 60S subunits (H). **I-P**) Eight cryo-EM maps corresponding to nutrient-deprived MCF-7 cell lysates comprise elongation states (I-L) similar to those in panels A-D; hibernating head-swiveled state with eEF2, CCDC124, pe/E tRNA and putative LARP1 (M), hibernating rotated ribosome with eEF2, SERBP1 and P/E tRNA (N); 60S with eEF2 and E-tRNA (O); 60S with eIF6 (P). (Q) Particle distributions among functional ribosome states in normal and nutrient-deprived MCF-7 cells. (R) Putative extended eEF1A conformation; the close-up box shows density with the extended eEF1A model fitted (green; PDB ID:8B6Z ^[22]^).

In lysates made from nutrient-deprived MCF-7 cells, the percentage of elongation-like ribosomes bound to mRNA and tRNAs decreased to 58% (Fig. 2I-L, Q), consistent with reduced translation. The abundance of unassociated 60S subunits increased from ∼10% to ∼15% (Fig. 2G, H, O, P, Q), in keeping with depletion in translation initiation. In both datasets, a fraction of 60S subunits is bound with elongation factor eEF2 (Fig. 2G, 2O). This is unexpected because the primary function of eEF2 is to translocate mRNA and tRNAs on fully assembled 80S ribosomes [53]. Indeed, eEF2 has been predominantly found on 80S ribosomes in this and previous studies. We discuss the implications of eEF2 binding to the 60S subunit in the next section.

Another notable difference between normal and nutrient-deprived MCF-7 cells concerns non-translating 80S ribosomes formed without an mRNA, also termed “hibernating” ribosomes [35, 54, 55]. Their content rises from 18% to 27% upon nutrient deprivation (Figs. 2F, 2M, 2N and 2Q). All hibernating ribosome classes feature eEF2. In addition, these maps contain Serpine1 mRNA-binding protein 1 (SERBP1) (Fig. 2F and 2N), suggested to be the predominant eukaryotic hibernation factor [55]. In nutrient-deprived cells, an additional hibernation class appeared, which contains coiled-coil-domain protein CCDC124 in the ribosomal P-site (Fig. 2M), resembling CCDC124-bound ribosomes purified from stressed cells [34] and in stressed ER-bound ribosomes [22]. Helical density in the mRNA tunnel in this class does not align with SERBP1 and likely corresponds to a different protein, such as La-related protein 1 (LARP1), as discussed below. Similarly, the non-translating 80S classes contain eEF2 in the BSC-1 dataset, with SERBP1 being the second most abundant hibernation factor with ∼32% of the total population (**Figure 2—figure supplement 1**).

To further investigate the potential functional roles of eEF2 on 60S subunits and hibernating 80S ribosomes, we obtained larger datasets of commercial RRLs, which contain larger fractions of non-translating 80S ribosomes.

### RRL contains non-translating ribosomes with eEF2 and SERBP1 or LARP1

2DTM with the 60S template, as described above, yielded ∼1.1 million particles in cryo-EM data collected from commercially available rabbit reticulocyte lysates (**Figure 3—figure supplement 2**). Such preparations are derived from nuclease-treated lysates to inhibit translation from endogenous mRNAs, allowing studies of reporter constructs [56]. Indeed, we collected data from translationally active lysates, as evidenced by luminescence from a NanoLuciferase reporter mRNA (Fig. 3A). After translating a reporter mRNA for 10 minutes, only ∼12.4% of ribosomes are bound with mRNA and tRNAs, indicating a depletion of translationally active states (Fig. 3 and **Figure 3—figure supplement 2**). This finding coincides with biochemical estimates of ∼10% of RRL ribosomes remaining on a reporter mRNA following a burst of translation [56]. The predominant elongating ribosomes feature eEF1A, A/T, P and E tRNAs (Fig. 3C) or eEF2 with chimeric ap/P and pe/E tRNAs resembling the nearly post-translocated state (POST-2: PDB ID 6GZ4; Fig. 3D). In addition, an unusual class emerged with density at the GTPase center that could not be assigned to canonical elongation factors (Fig. 3F, **Figure 3—figure supplement 3B**). The shape of the density most closely resembles developmentally regulated GTP-binding proteins DRG1 or DRG2, whose homologs (Rbg1/2) were recently visualized on yeast ribosomes [57, 58]. The protein interacts with the A-site tRNA—as if stabilizing the acceptor arm docked at the peptidyl transferase center—consistent with the proposed role of DRG and Rbg proteins in restoring elongation on stalling-prone sequences, such as poly-proline [57, 59]. Whereas the overall shape and secondary structure resemble DRG1 or DRG2, the local resolution is insufficient to distinguish between these or other similarly structured proteins. Both yeast and mammalian counterparts are reported to function with a companion factor (Tma146p or Gir2 in yeast; or DFRP1 and DFRP2 in mammals), but our maps do not contain density that could correspond to DFRP1/2 near the putative DRG1/2 density. Future work will elucidate the function of these or other DRG-like GTPases in the context of an elongation complex.

**Figure 3.**
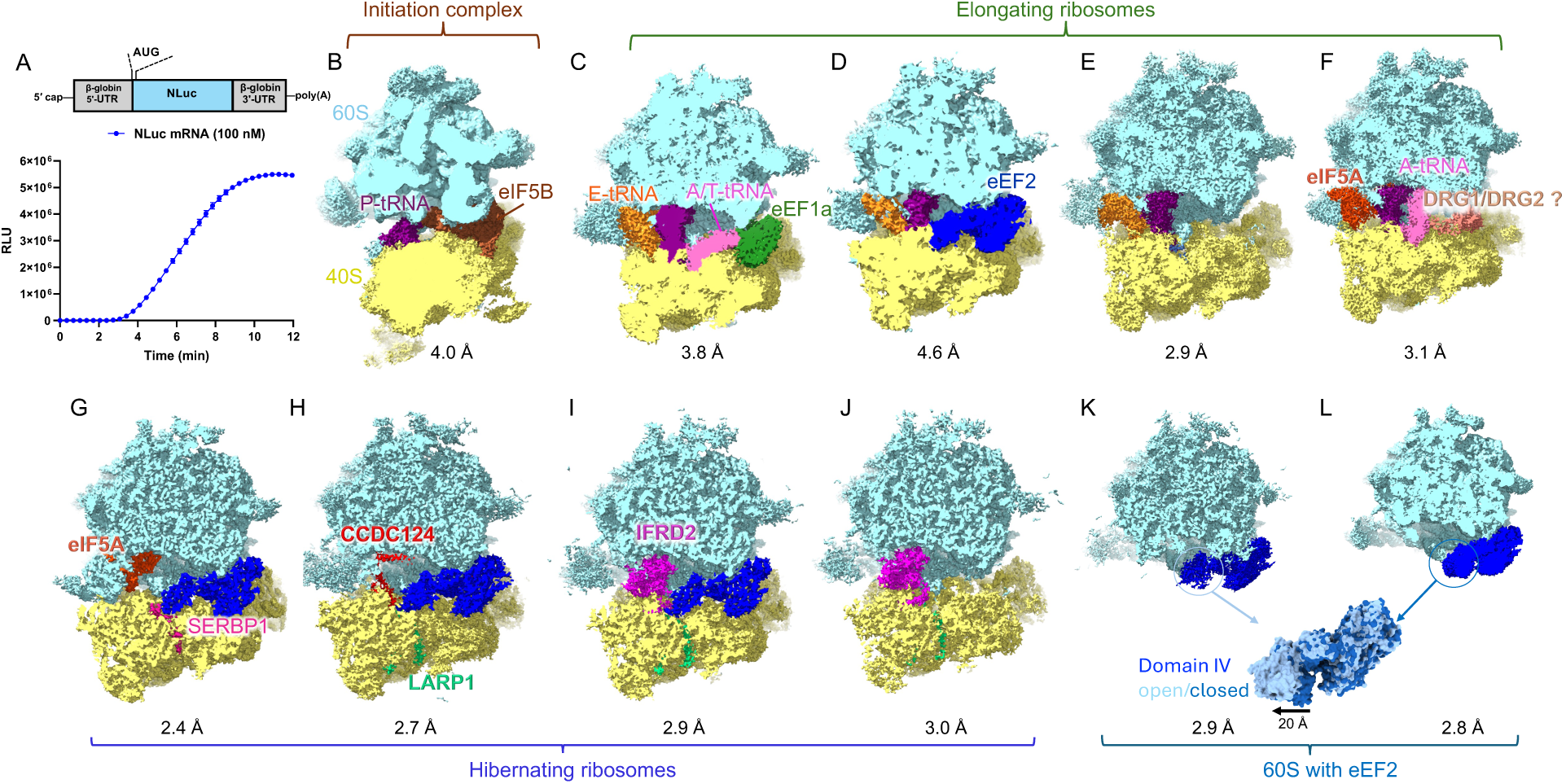
*In extracto* cryo-EM of rabbit reticulocyte lysates. **A**) NLuc mRNA construct used to monitor NanoLuciferase translation (top) and real-time translation kinetics (bottom). The graph represents the mean ± SEM from three independent experiments. **B-L)** Cryo-EM maps of the major classes resulting from 3D classification: (B) initiation complex with eIF5B; C-F) elongating ribosomes: codon-sampling with eEF1A (C), post-translocation with eEF2 ap/P and pe/E tRNAs (D), pre-translocation hybrid-state (E), and with GTPase density putatively assigned to DRG1/DRG2 next to A/A tRNA (F). G-J) Hibernating ribosomes: 40S-rotated ribosome with eEF2, eIF5A, and SERBP1 (G); 40S-head-swiveled ribosome with eEF2, CCDC124, and LARP1 in the mRNA tunnel (H); 40S head-swiveled with eEF2, IFRD2, eEF2 and LARP1 in the mRNA tunnel (I); 40S-head-swiveled with IFRD2 and LARP1 (J); 60S subunit with eEF2 domain IV open (K); 60S with eEF2 domain IV closed (L): the extent of domain IV movement in 60S-bound eEF2.

Most ribosome particles in RRL—nearly 60%—are in non-translating states, evidenced by the absence of mRNA density. Like in primate lysates described above, eEF2 is the predominant translation factor bound to 97% of hibernating ribosomes (Fig. 3). The most abundant class represents nearly half of hibernating ribosomes and adopts a partially rotated ribosome conformation. Here, the small subunit body is rotated by ∼5° and the head is swiveled by ∼20° relative to a non-rotated decoding state (5LZS; Table S3), so that the ribosome resembles a late translocating ribosome with two “chimeric” ap/P and pe/R tRNAs and eEF2 or EF-G [60]. The most abundant hibernating ribosome, however, features no tRNA, and instead three proteins overlap with the three tRNA binding sites: eEF2 spanning from the sarcin-ricin loop (SRL) on the large subunit to the decoding center (DC) on the small subunit; SERBP1 occupying the mRNA tunnel extending into the P site; and eIF5A extending from the E site into the peptidyl transferase center, PTC (Fig. 3G). As such, the ribosomal RNA at the key functional centers—SRL, DC and PTC—are inaccessible to other proteins, which was proposed to serve as protection against nucleolytic cleavage that may occur in stressed cells [61–64].

Our classification also yielded three previously unreported, to our knowledge, states of hibernating ribosomes that adopt non-rotated conformations (body rotation less than 1°), in which the head of the small subunit is swiveled by ∼18° (Fig 3. H-J). Like in the predominant state described above, the ribosomal functional centers are protected by the simultaneous binding of eEF2 with CCDC124 (Fig. 3H) or eEF2 with IFRD2 (interferon-related developmental regulator 2; Fig. 3I). Remarkably, helical density in the mRNA tunnels in these maps differs from the coiling SERBP1 (Fig. 4) and most closely resembles La-related protein 1 (LARP1; residues 659-724), which was recently imaged by cryo-EM on vacant 40S ribosomes [32]. LARP1 is a component of RRL [65] and is implicated in translation initiation on pyrimidine-rich mRNAs [66]. LARP1 was shown to bind 80S ribosomes [32], although no structure was reported.

**Figure 4.**
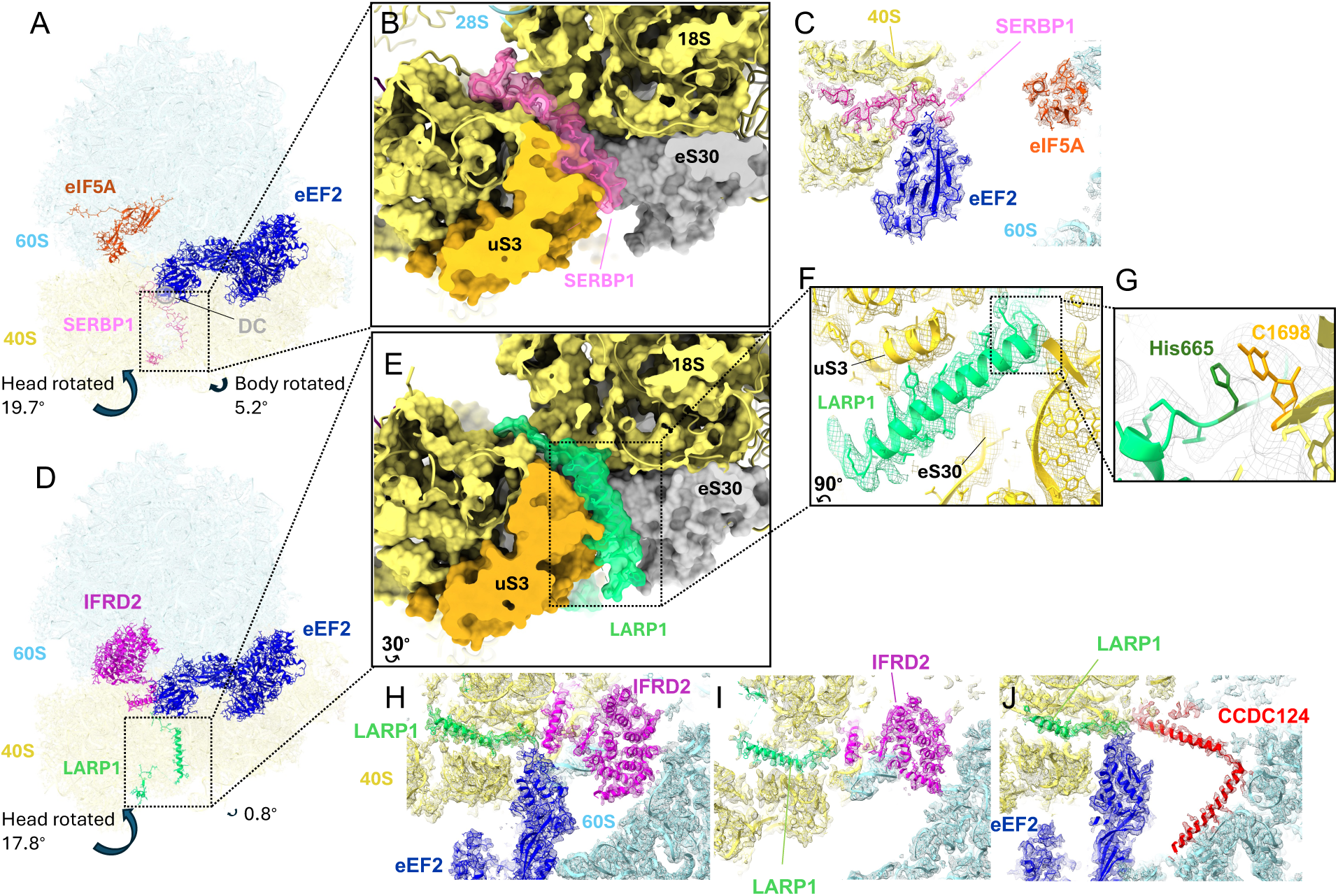
The occupancy of the mRNA tunnel in hibernating ribosomes from RRL. **A**) The 80S ribosome with eEF2, eIF5A and SERBP1 (DC: decoding center). (**B**) Close-up view of cryo-EM density for SERBP1 in the mRNA tunnel. (**C**) Density for SERBP1 interaction with eEF2. (**D**) The 80S ribosome with eEF2, IFRD2 and LARP1. (**E**) Close-up view of LARP1 density in the mRNA tunnel. **F-G**) Views of the density for the LARP1 helix interacting with 40S ribosomal proteins and RNA. **H-J**) Similar densities for LARP1 in CCDC124- and IFRD2-bound ribosomes.

Our finding suggests that instead of SERBP1, LARP1 or another protein with LARP1-like binding mode binds the hibernating 80S ribosomes adopting a head-swiveled state in the presence of eEF2 with CCDC124 or IFRD2. Due to density features that closely resemble LARP1 (Fig. 4) and LARP1 regulation during stress (see below), we hypothesize that the density corresponds to a LARP1 isoform. The protein interacts simultaneously with eEF2 and CCDC124 or IFRD2 in the decoding center (Fig. 4), and this interaction likely accounts for the ribosome rotational state. The functional interplay between CCDC124 and LARP1, as well as between IFRD2 and LARP1, is intriguing. Their simultaneous recruitment to the hibernating ribosomes may enhance their individual roles as translational repressors during stress and may be cell- or tissue-specific. In addition, as LARP1 controls translation of mRNAs with a terminal oligopyrimidine (TOP) motif, which encode ribosomal proteins, the protein’s recruitment into hibernating ribosomes may help downregulate ribosome expression during stress. Finally, the binding of the extra-ribosomal DM15 domain of LAPR1 to TOP mRNAs may prepare these mRNAs for translation upon stress relief. Indeed, like SERBP1, LARP1 is the substrate of TORC1 [67]. TORC1 was shown to stimulate translation at least in part by activating dormant ribosomes via phosphorylation of Stm1, the yeast counterpart of SERBP1 [68]. LARP1 binding to and release from 80S ribosomes may similarly be controlled by TORC1 and Akt/S6K1[67] to switch between stress and proliferation conditions.

The hibernating ribosomes imaged directly in RRL lysates differ from ribosomes purified from RRL [35], in that the latter only contained eEF2 with SERBP1, or tRNA with or without IFRD2, but not eIF5A, or eEF2 with IFRD2/CCDC124, and LARP1. While some of these proteins may be low in abundance and may have fallen below the detection limit of the previous study, the absence of highly abundant eIF5A and eEF2 in the corresponding classes indicates that they likely dissociated from ribosomes during purification. Indeed, fractions of IFRD2-bound ribosomes with or without eEF2 in the RRL lysate data (Figs. 3I-J) suggest transient eEF2 interactions with some types of hibernating ribosomes. An internal loop of IFRD2 (aa 400-409), ordered in the absence of eEF2, is displaced by the tip of eEF2 domain IV, suggesting that steric hindrance may account for the partial occupancy of eEF2 in IFRD2-bound ribosomes (**Figure 3—figure supplement 5**). Nevertheless, the most abundant class with eEF2, eIF5A, and SERBP1 is readily detected in cellular cryo-EM or single-particle cryo-EM studies from other mammalian cell lines [69], whereas no CCDC124- or IFRD2-bound ribosomes have been reported with LARP1. These findings underscore the importance of imaging ribosomes within cells or lysates to enable the identification of cellular interactions.

The 60S classes contain eEF2•GDP bound next to the sarcin-ricin loop, featuring open (Fig. 3J) and compact (Fig. 3K) conformations. eEF2 and its bacterial counterpart EF-G rearrange between open and compact conformations on the complete translocating ribosome [12, 70] or when free in solution [71]. Capturing these conformations in the 60S subunit likely reflects an equilibrium that eEF2 is spontaneously sampling, resembling that of free EF-G [72]. A potential role of 60S•eEF2 complexes may be to preserve the SRL in free subunits under stress.

In addition, 60S classes feature density spanning the polypeptide tunnel (**Figure 3—figure supplement 4A-B**), resembling 60S maturation factor ZNF622 in mammals [73] and its homolog Rei1 in yeast [74]. The density differs from both factors on the 60S periphery, where it approaches EBP1 (**Figure 3—figure supplement 4C-D**), suggesting a different conformational state of ZNF622 or another protein, whose role may be to occlude the PTC and tunnel periphery during 60S hibernation. Future studies will bring insights into the roles of the protein(s) and into the functions and transitions of 60S•eEF2 complexes to the pool of translating ribosomes.

### Structural basis for eEF2•GDP binding to hibernating ribosomes

The essential cellular function of eEF2, and of its better studied bacterial counterpart EF-G, is to catalyze translocation of mRNA and tRNA during elongation [10, 75–78]. To this end, the translocase binds to a pre-translocation ribosome, containing peptidyl-tRNA in the hybrid A/P state and deacyl-tRNA in the P/E state (in the tRNA hybrid-state nomenclature, the first letter denotes the position on the small subunit and the second letter on the large subunit). The ribosome adopts a rotated state, in which the small subunit is rotated by ∼10° relative to its position in a non-rotated ribosome harboring the “classical-state” A and P tRNAs (aka A/A and P/P). Upon binding, the GTPase domain of eEF2/EF-G docks at the universally conserved sarcin-ricin loop (SRL) critical for triggering GTP hydrolysis [79–82]. The translocase domain (domain IV) is placed next to the A site on the small subunit. Due to the ribosome’s inherent propensity to undergo subunit rotation [83], the spontaneous reversal of the small subunit results in the arrival of translocase domain IV to the A site (also known as the decoding center) and the movement of tRNA-mRNA helix from the A to P site on the small subunit [84, 85]. The enzyme dissociates upon the completion of subunit rotation and translocation after GTP hydrolysis and P_i_ release from the GTPase domain, resulting in the departure of this domain from SRL [12].

While the roles of eEF2 and GTP hydrolysis in tRNA-mRNA translocation are reasonably well understood, their functions in ribosome hibernation remain a puzzle. Unlike the preferred translocase substrate—the pre-translocation ribosome—hibernating ribosomes do not contain peptidyl-tRNA or mRNA in the A site, which could stabilize eEF2 on the ribosome. Furthermore, previous studies captured eEF2 on hibernating ribosomes in the presence of SERBP1 in the decoding center, suggesting that the latter might contribute to eEF2 stabilization on the ribosome [55]. Indeed, SERBP1, spanning the mRNA tunnel, contacts domain IV of eEF2 in the A site in this (Fig. 4A-C) and previous studies [86]. Our work, however, also identifies eEF2 on hibernating ribosomes in the presence of several other factors, or in their absence (on the 60S subunit), suggesting novel insights into the function of eEF2 on hibernating ribosomes.

In hibernating 80S ribosomes, eEF2 interacts with SERBP1 or LARP1 (Fig. 4), arguing that the latter can replace SERBP1 in its function to stabilize eEF2 in the A site of the small subunit. By contrast, eEF2 binding to free 60S in this work, as well as binding of eEF2/EF-G to rotated ribosomes with a single tRNA [35, 45, 82] placed away from domain IV, argue that the contact in the 40S A site is not necessary for eEF2 binding. Instead, binding on the 60S subunit may be the predominant interaction to stabilize eEF2.

eEF2 binds to the hibernating ribosomes or 60S subunits similarly to that in translocation complexes, in which the GTPase domain is placed next to the SRL (Fig. 5). The binding of eEF2/EF-G to pre-translocation ribosomes requires a GTP-bound conformation (e.g. GTP, non-hydrolysable GTP analogs [87, 88], or GDP+Pi [12]), as the ordered GTPase switch loops bridge the large subunit with the rotated small subunit [12]. On translocating ribosomes, the release of Pi after GTP hydrolysis and subunit rotation is coupled with switch-loop disordering and translocase dissociation [12]. Remarkably, we find that hibernating 80S ribosomes, in both rotated and non-rotated conformations, feature eEF2 with GDP (Fig. 5a), and that the eEF2 switch loops remain ordered on the rotated ribosome.

**Figure 5.**
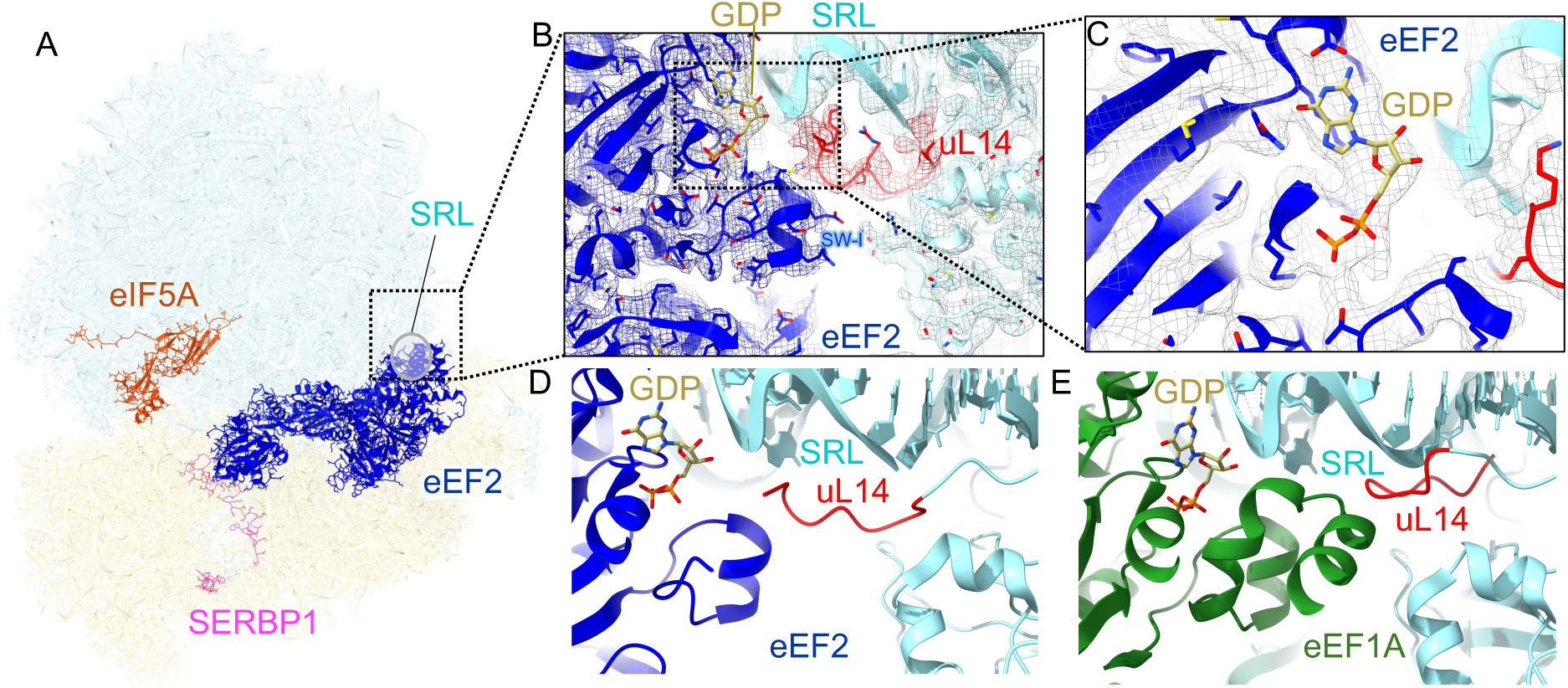
Interactions of eEF2 with the GTPase-activating center in hibernating ribosomes. **A**) Overall view of the 80S structure with eEF2, eIF5A, and SERBP1. **B**) Cryo-EM density and model of the eEF2 GTPase center at the SRL. **C**) GDP density in the GTPase center of the predominant hibernating ribosomes. **D, E**) Distinct conformations of the N-terminal tail of uL14, highlighted in red, in ribosomes bound with eEF2•eIF5A•SERBP1 (D) and with eEF1A•GDP (PDB ID, 5LZS) (E).

Our examination of the eEF2-containing cryo-EM maps revealed that the GTPase domain is held at the SRL near ribosomal protein uL14. The N-terminal tail of uL14 has been unmodeled in previous structures of mammalian ribosomes with GTPases, likely due to disorder. Indeed, in cryo-EM maps of ribosomes bound with eukaryotic elongation factor 1A (eEF1A) [89] or release factor 3 (eRF3) [89], harboring homologous GTPase centers, density for the first 10 residues of uL14 is absent or placed farther from the GTPase centers. By contrast, continuous density is evident in our maps of eEF2-bound ribosomes and indicates that the N-terminus is reaching within 5 Å of GDP in the GTPase center of eEF2 (Fig. 5B). The switch-I (SW-I) helix of eEF2 (aa 55-66) contacts residues 4-7 of uL14 (Fig. 5B). This interaction resembles that in recent structures of *G.gallus* eEF2 bound to translocation-like ribosomes [90], although the path of the N-terminal tails of uL14 was modeled differently (**Figure 4—figure supplement 1**). Similarly, continuous density is observed for the N-terminal tail of uL14 in our 60S•eEF2 complexes, where domain IV also contacts helix 69 of 28S rRNA (**Figure 4—figure supplement 2**), suggesting additional stabilization of eEF2. In sum, our findings demonstrate that SERBP1 is not required to stabilize eEF2 in the absence of mRNA and tRNA. The N-terminus of uL14, which adopts different conformations in the presence of different translational GTPases, may be involved in stabilizing eEF2 on the ribosome or isolated 60S subunits.

### Conclusions

We demonstrate that *in extracto* cryo-EM enables high-resolution visualization of ribosome complexes in the presence of cellular components. In comparison with canonical single-particle cryo-EM (usually involving purified macromolecules) and cellular cryo-EM/ET, this method offers advantages of both methods (Table 1). First, the preservation of cytoplasmic components in lysates enables the identification of interactions between cytoplasmic macromolecules. Indeed, the identification of eEF2 and other novel factors in 60S and 80S complexes in our work contrasts with approaches in which ribosomes were purified prior to cryo-EM analyses. Furthermore, the comparison of fresh lysates prepared from normally treated MCF-7 cells and nutrient-deprived cells demonstrates the reduction of actively translating ribosomes and increase in non-translating ribosomes and individual 60S subunits. Additional optimization of buffer conditions may be required to more accurately represent the translation states observed in cells, as ionic conditions are known to affect the conformation of the ribosomes (e.g. rotated/non-rotated) and binding of protein factors [91–94]. For cells or samples with lower abundances of ribosomes or other macromolecules/complexes of interest, a lysate concentration step or collection of a larger dataset may be considered. Nevertheless, our initial analyses demonstrate the expected changes in the translational machinery, consistent with translational repression during nutrient deprivation in cells [95–98].

**Table 1.**
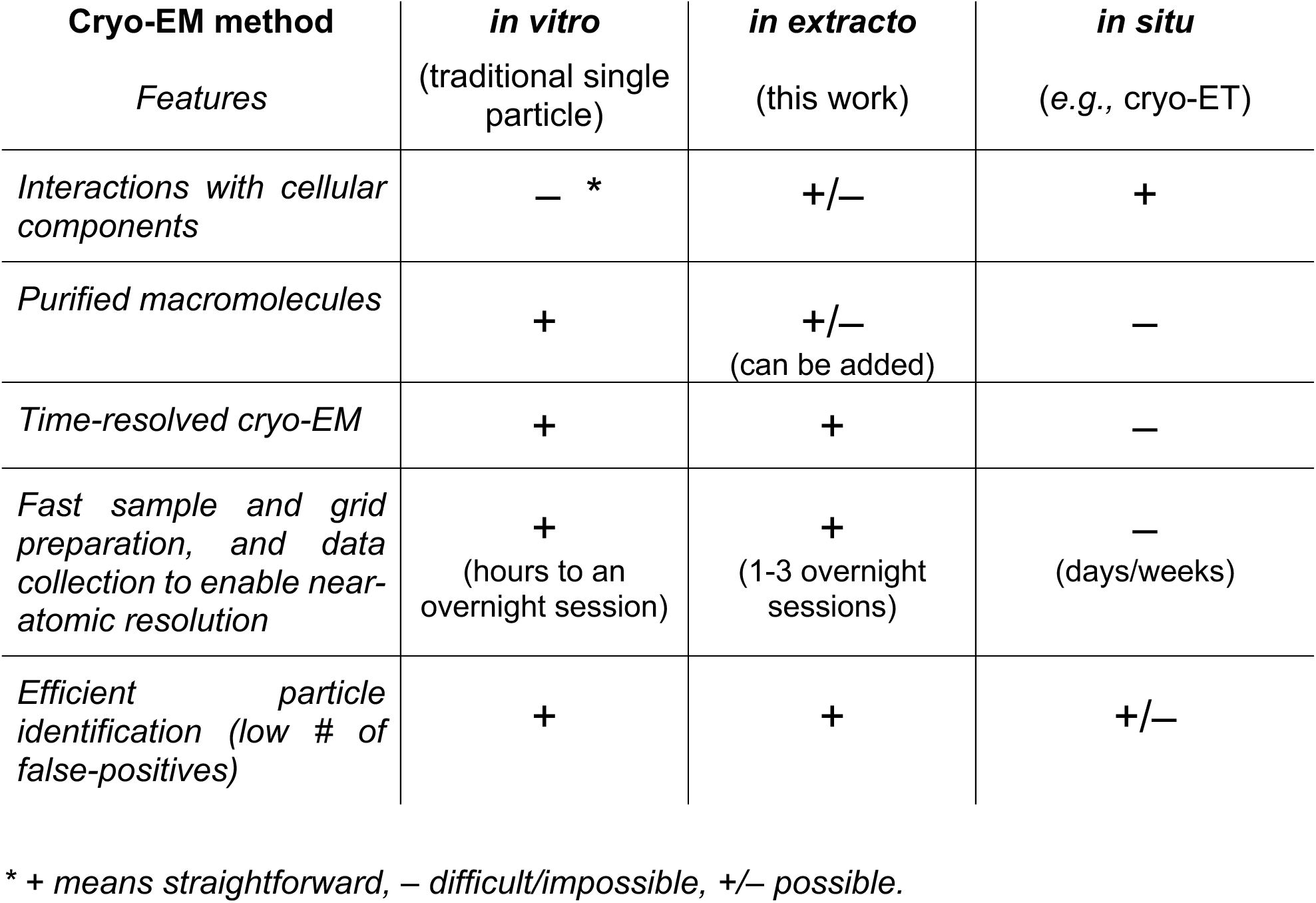
Comparison of *in extracto* cryo-EM with traditional *in vitro* and *in situ* cryo-ET approaches to structural biology.

*In extracto* cryo-EM is a flexible method allowing to add purified components, such as an mRNA reporter (Fig. 3), to lysates at different concentrations and time points, enabling equilibrium and time-resolved cryo-EM studies, resembling *in vitro* single-particle cryo-EM [12, 14, 36]. Although purified components could also be added to cells, their intracellular concentrations and time course are difficult to control and measure, making conditions in *in situ* cryo-EM more challenging to control than in *in extracto* cryo-EM. While this manuscript was in preparation, canonical single-particle cryo-EM analyses of bacterial lysates were reported, revealing numerous 70S ribosome structures [99]. Similarly to mammalian lysates, bacterial systems are widely used with different reporters and translation factor combinations [100, 101], further underscoring the flexibility and broad applicability of *in extracto* cryo-EM.

Finally, we demonstrate that fast grid preparation with lysates (∼10 minutes) enables data collection similar to or even faster than traditional single-particle cryo-EM requiring sample purification. The resulting sample layers are thicker than those of purified macromolecules, necessitating the use of template matching, such as 2DTM and GisSPA [102], which are slower than conventional particle picking methods implemented in most cryo-EM software packages. Yet, since no FIB milling of cellular samples is necessary for lysates, the overall throughput including sample preparation and fast GPU-accelerated data processing is superior to *in situ* cryo-ET or cryo-EM. Furthermore, although 2DTM may be computationally more expensive than traditional template-based particle picking employed with *in situ* samples, the latter is often subject to a large number of false positives [103] that have to be removed in subsequent image processing steps.

In this work, we use cryo-EM to grant a unique glimpse into the translation system that has been used in hundreds of studies in the past five decades [104, 105]. In comparison with freshly made lysates from MCF-7 cells featuring ∼55-70% ribosomes bound to an mRNA (Fig. 2), the translationally active commercial nuclease-treated RRL contains a much smaller ∼12% fraction of elongating ribosomes following the translation of an added mRNA construct. These findings are consistent with previous estimates of mRNA-bound ribosomes [30] and likely indicate the expedient exhaustion of the added resources, degradation of mRNA and/or other time-sensitive aspects of the cell-free lysate.

Nevertheless, thanks to the large fraction of non-translating ribosomes in RRL, our analyses have substantially expanded the understanding of the translational hibernation repertoire. We find that eEF2 is bound to the majority of hibernating ribosomes, highlighting that eEF2 functions beyond its canonical role of the translocase. The binding of eEF2 to hibernating 80S and 60S can occur in the absence of the previously identified abundant hibernation factor SERBP1, underscoring that the stabilization by SERBP1 is not required for this binding. Instead, eEF2 appears to be stabilized predominantly by the interactions with the SRL region on the large subunit. These interactions feature the N-terminus of uL14 approaching the GDP molecule in the eEF2 active site. The conformation of the N-terminal uL14 tail differs from those in eEF1- and eRF3-bound complexes, suggesting that the tail may contribute to stabilizing eEF2.

We also find that LARP1 is an abundant hibernation factor occupying the mRNA tunnel in 80S ribosomes. As discussed above, the simultaneous recruitment of LARP1 with IFRD2 or CCDC124, two other translational repressors, may serve to both protect all ribosome active centers and enhance their translational repression potential. Furthermore, LARP1 may co-localize TOP mRNAs with hibernating ribosomes to expediently resume translation of ribosomal proteins when stress subsides.

In summary, although the composition of hibernation factors differs among hibernating ribosomes, their binding to the SRL (eEF2), mRNA tunnel and decoding center (SERBP1 or LARP1), P-site and peptidyl-transferase center (eIF5A, CCDC124 or IFRD2) make these sites inaccessible for nucleolytic or other enzymes (Fig. 6). These strategic positions are consistent with the proposed function of hibernating ribosome complexes being kept “in reserve” to enter the pool of translating ribosomes when necessary [106, 107]. Together, our findings underscore the opportunities to uncover novel cellular interactions using *in extracto* cryo-EM.

**Figure 6.**
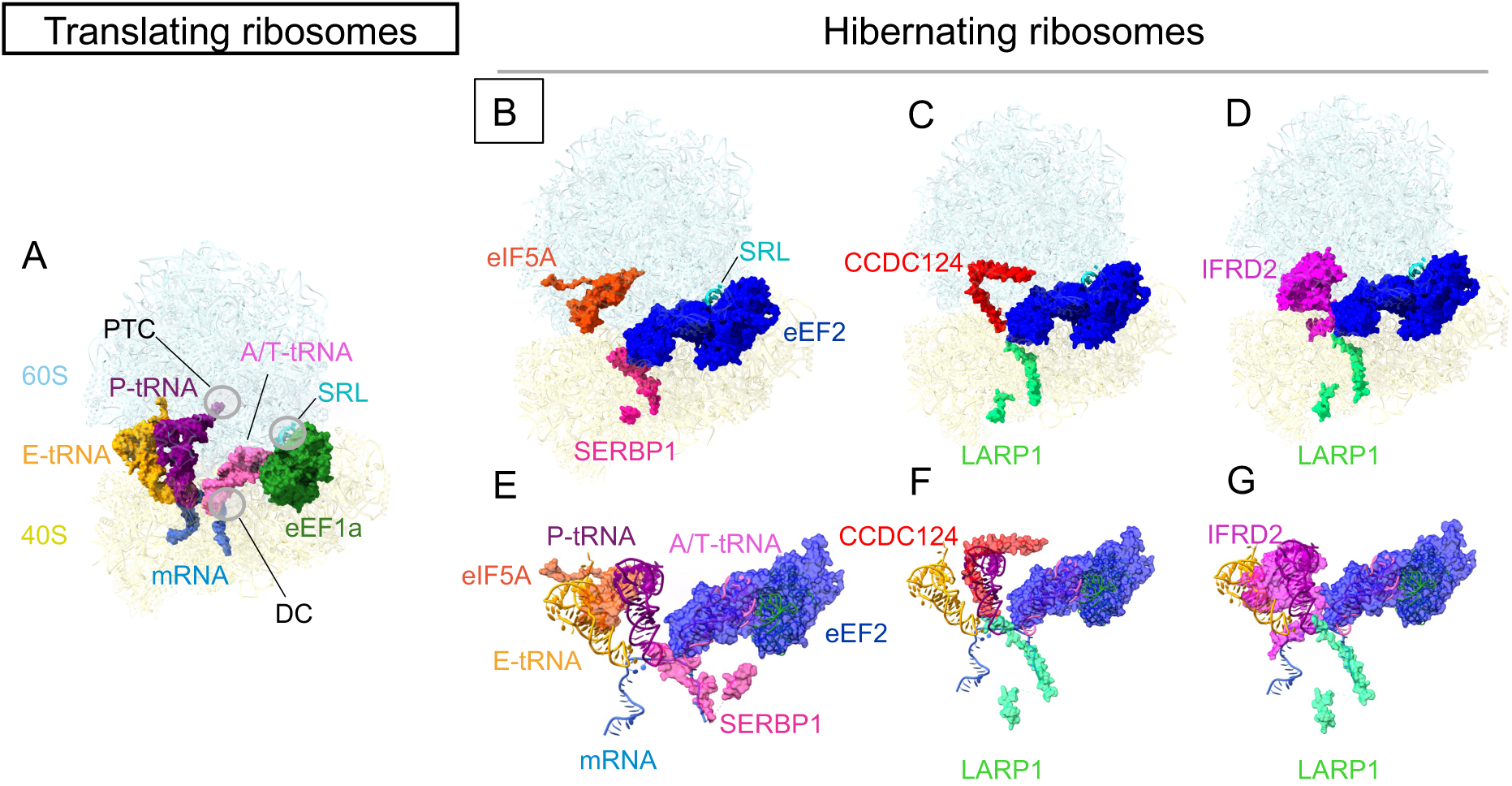
Binding sites of hibernation factors overlap with ribosomal functional centers. **A-D**) Comparison of a translating ribosome (PDB 5LZS and mRNA from PDB 4V6F; panel A) with hibernating ribosomes identified in this work. **E-G**) Superposition of translating and hibernating ribosomes illustrates that the key ribosomal functional centers all shielded by hibernation factors (eEF1A, A/T-tRNA, P-tRNA and E-tRNA are from PDB 5LZS and mRNA from PDB 4V6F). Panel E shows the view from panels B-D rotated by 36°.

## Methods

### Optimization of 2DTM in RRL data

To study angiogenin interactions with translating ribosomes, we collected a test cryo-EM dataset of translationally competent commercial lysate supplemented with recombinant angiogenin as described [36]. The sample contained 33% rabbit reticulocyte lysate (without nuclease treatment; Green Hectares), 1 µM angiogenin, and 2.5 ng/µl nanoluciferase mRNA and was applied to grids and vitrified using standard single-particle cryo-EM conditions [36]. During grid screening, we readily distinguished ribosomes in thin ice areas of the grid and performed overnight collection yielding 10,084 micrographs [36]. Standard single-particle cryo-EM pipelines, however, either failed to yield high resolution reconstructions or discarded most of the particle data (Table S1). We first utilized our standard pipeline aligning movies in IMOD [108], importing micrographs into *cis*TEM for CTF-correction and particle picking (272,478 particles), and searching alignment angles at low resolution (30-60 Å) against a 25-Å low-pass-filtered rabbit ribosome map (EMDB 4729) in Frealign 9.11 [109]. The reconstructions displayed low resolution (>30 Å) and were dominated by a preferred orientation. Limiting the micrographs to ones where CTF Thon rings could be fitted to better than 3.2 Å resolution (600 micrographs with 7,915 particles) and manual particle picking (removing areas of thick carbon or crystalline ice, recentering some picks) were required to accumulate a 6,126-particle stack, yielding a reconstruction at 3.1 Å resolution (FSC 0.143). The cryo-EM map was further classified into 6 or 8 classes yielding four types of classes: 80S+Angiogenin+strong Ternary Complex (80S+Ang+strong TC), 80S+Angiogenin+weak Ternary Complex (80S+Ang+weak TC), hibernating 80S with eEF2, and 60S at 3.4-4.2 Å resolution. We also tried to use a Relion v3 [110] pipeline with CTF correction in GCTF, particle picking using the Laplacian-of-Gaussian picker (140,721 particles), with two rounds of sorting and 8 rounds of 2D classification to remove ice contaminants and particles too small to be ribosomes, then aligning picked particles to a reference derived from EMD-4729, yielding an initial reconstruction at lower than 6.64 Å resolution (4× binned particle stack) from 52,580 particles. Two rounds of 3D classification yielded the same three classes as the *cis*TEM pipeline and at similar resolution (up to 3.2 Å) from ∼16,480 particles. Thus, despite imaging in cell lysates, with manual particle picking we could distinguish and align some ribosome particles and find a novel, high-resolution state (80S+Ang+TC).

Inspection of the micrographs suggested that many ribosome particles were missed by the manual picking process. We therefore employed particle picking with 2DTM [21]. We started with a limited particle stack of the 1,600 best micrographs from a total of 10,084 micrograph total and picked with a high-resolution template prepared from the 60S of the 80S•poly(GR) complex (PDB:7TOR [111]). The coordinates corresponding to the 60S subunit were saved as a separate PDB and converted into a .mrc map using the e2pdb2mrc.py program from EMAN2 [112] using a super-sampled pixel size of 0.415 Å, box size of 1024, and specifying that coordinates should be centered. The map was resampled to 2× using resample.exe from cisTEM [21] and then B-factor filtered using bfactor.exe from Frealign 9.11 [109], applying a B-factor of 80 Å^2^ as described previously [21]. Particle picking with 2DTM in cisTEM yielded a stack of 37,859 particles, of which >95% belonged to high-resolution classes. 3D classification of this larger stack revealed states that were not previously separated in the *cis*TEM/Frealign v9.11 nor Relion 3 pipelines.

We optimized high-resolution template matching procedures for faster performance and maximum particle number with high signal/noise ratio. To this end, we performed a grid search for out-of-plane angles, in-plane angles, and defocus (z direction) search parameters on a limited set of micrographs. We found that increasing default values of out-of-plane angles from 2.5 to 3.5 and in-plane search angles from 1.5 to 2.5, omitting the defocus search, and resampling the micrographs and the search template to a 1.5 Å/pixel optimally speeds up the procedure without substantially sacrificing the number of picked particles. Using these parameters, we picked the full set of micrographs, excluding off-target grid locations (black images, images with broken ice, images on thick carbon), with the 60S template yielding 88,488 particles. The new dataset was rich in substates, including collided ribosomes in the translationally competent 80S•eEF2 and different positions or identities of ternary complexes in 80S•Ang•TC (see Table S1). Thus, high-resolution template matching identifies ribosome particles in micrographs from cell lysates and reveals an abundance of novel states for analysis as compared to traditional single-molecule processing from select micrographs.

We found it beneficial to collect data from grid regions with thicker ice, where the population of ribosomes is higher than in thinner-ice regions of the grid, which contrasts the strategies for conventional single-particle grid and data optimization. To this end, we optimized the use of defocus search for each dataset or a batch of micrographs. We first grouped micrographs based on their CTF fit resolutions and selected ten micrographs representing the highest and lowest fit resolutions. Template matching was then performed with the defocus search enabled or disabled, and the number of picked particles for each micrograph was compared. When a meaningful difference (in some, cases, two-fold or more) was observed—commonly for thicker-ice micrographs—we enabled defocus search for the corresponding groups of images. This approach allows to maximize the dataset size to enable extensive classification and improved resolutions of the resulting reconstructions.

### Preparation of MCF-7 and BSC-1 cell extracts

Cell extracts were prepared from MCF-7 cells by using digitonin buffer extraction, as in previous studies, with modifications [42, 113, 114]. MCF-7 cells (obtained from Dr. Michael Green’s lab [115]) were cultured in DMEM (Gibco) supplemented with 20% FBS (Gibco), and 10% of Penicillin-Streptomycin (Gibco). To prepare non-stressed cells, MCF-7 cells were seeded in 75 cm² flasks, with the media refreshed every 24 hours and 6 hours before cell collection, for 48 hours. For cells with nutrition stress, the media was not changed within 48 hours. Four flasks of cells were used for lysate preparation with a total of 8 ml of semi-permeabilization buffer (25 mM HEPES pH 7.2, 110 mM KOAC, 15 mM Mg(OAC)2, 1mM Dithiothreitol (DTT), 0.015% Digitonin, 2x Protease Inhibitor (Roche), 40 U/ml RNase-In (Promega), 1 mM EGTA). Cells were washed twice with ice-cold PBS buffer (Gibco) and then 1 ml of semi-permeabilization buffer was added to each flask. After a 5-minute incubation with the semi-permeabilization buffer, the cytosol-containing mixture was collected. This step was repeated to maximize sample collection (all this procedure was done in the cold room at 4°C). 8 ml of sample were transferred to 30 kDa MWCO Amicon filters and concentrated until the RNA concentration (A_260_; NanoDrop One Spectrophotometer, Thermo Scientific) exceeded 1000 ng/µL. Concentrated cell extract was directly added to the grids and vitrified as described below (Plunge freezing). To skip the concentration step, 200 μl of semi-permeabilization buffer (instead of 1 ml) can be added to each flask and collected for direct application to a grid.

BSC-1 cells (ATCC) were cultured in DMEM (Invitrogen) supplemented with 10% HI-FBS, 2 mM GlutaMAX (Gibco) and 50 ug/mL of Penicillin-Streptomycin (Gibco). Cells were seeded in a 75 cm² flask and cultured to confluence. After being washed twice with pre-warmed PBS, the cells were detached with Trypsin-EDTA (Gibco) and collected by centrifugation at 300 × g for 4 minutes. The cell pellet was resuspended in 100 μL of semi-permeabilization buffer (25 mM HEPES, 110 mM KOAc, 15 mM Mg (OAc)_2_, 1 mM DTT, 0.015% digitonin, 2× protease inhibitor cocktail (Roche), 40 U/mL RNaseIn (Promega), 1 mM EGTA) at 4°C for 5 minutes. Following centrifugation at 1,000 × g for 5 minutes, the supernatant was collected for use in grid preparation (see below). RNA concentration was quantified prior to grid preparation as a quality control measure, with no dilution of the extract performed.

### Preparation of RRL samples for cryo-EM

Commercial micrococcal-nuclease-treated rabbit reticulocyte lysates (Promega; L4960) were supplemented with the following list of buffer and reagents to reach the final concentration of 50% RRL, 30mM HEPES-KOH, pH7.5, 50mM KOAc, 1 mM Mg(OAc)_2,_ 0.2 mM rATP, 0.2 mM GTP, 0.02 mM Amino acid mix minus Met (Promega), 0.02 mM Amino acid mix minus Cys (Promega), 5mM DDT added with MilliQ-Water. For grid preparation, two samples of RRL were prepared: 1) with mRNA encoding NanoLuciferase (see below); and 2) without mRNA. After thawing, 50% RRL (final concentration, prepared according to vendor instructions) was incubated for 5 min at 4°C. Then solution of NLuc mRNA (final concentration 25 ng/uL or ∼100nM, in water) or an equivalent volume of water was added and incubated for 10 min at 30°C. The mixture was directly applied to the grids and vitrified (see Plunge Freezing).

### Preparation of *in vitro* transcribed NanoLuciferase mRNAs

A plasmid carrying the coding sequence for NanoLuciferase flanked by the 5′- and 3′-UTRs of rabbit *HBB2* was synthesized by Azenta (vector: pUC-GW-Kan), as designed and described [56]. DNA templates for *in vitro* transcription were PCR-amplified from the plasmid using Phusion High-Fidelity DNA Polymerase (NEB; M0530L) and primers (Integrated DNA Technologies) containing the T7 promoter (5′-TTTTTTAATACGACTCACTATAGGGAGAACACTTGCTTTTGACACAACTGTG-3′) and a 30-nt poly-A-tail (5′-TTTTTTTTTTTTTTTTTTTTTTTTTTTTTTGCAATGAAAATAAATTTCCTTTATTAGCC-3′). After PCR, DNA templates were purified by phenol/chloroform extraction and dissolved in nucleases-free Milli-Q water. DNA concentrations were measured using a NanoDrop One Spectrophotometer (Thermo Scientific). The presence of a single band as DNA template was confirmed by agarose gel electrophoresis (1% (w/v) agarose in TAE buffer). *In vitro* transcription reactions were carried out using 4 μg of purified DNA templates and purified recombinant T7 polymerase in transcription buffer (166 mM HEPES–KOH, pH 7.5; 20 mM MgCl_2_; 40 mM DTT, 2 mM spermidine, 25 mM each of ATP, GTP, CTP, and UTP; and 40 U/μl RNase Inhibitor (NEB; M0314S)) in an 80 μl reaction. After incubation at 37°C for 3.5 h, magnesium pyrophosphate precipitate was removed by centrifugation (14,000 x g, 5 min), and mRNA was precipitated from the supernatant by adding LiCl (2.5 M final concentration) and incubating at –80°C overnight. The next day, mRNA was pelleted by centrifugation (21,300 x g, 15 min at 4°C), washed with cold 80% ethanol and pelleted again. This washing step was repeated three times. After discarding the supernatant, the mRNA pellet was air-dried and dissolved in nucleases-free Milli-Q water. To attach 5′cap, capping reactions were performed using the Vaccinia Capping System (NEB; M2080S) following the protocol. The 5′ capped mRNA was then purified by LiCl precipitation as described above and dissolved in nucleases-free Milli-Q water. mRNA concentration was determined from A_260_ absorbance using a NanoDrop One Spectrophotometer (Thermo Scientific). The size and integrity of the *in vitro* transcribed mRNA were examined by denaturing agarose gel electrophoresis (1% (w/v) agarose in MOPS buffer with 1.11% (v/v) formaldehyde) alongside an ssRNA ladder (NEB; N0362S). The stock solution was stored at −80 °C.

### *In vitro* translation in rabbit reticulocyte lysates

*In vitro* translation was performed using a commercial micrococcal-nuclease-treated RRL (Promega; L4960) with modifications as described below. Translation reactions were carried out in the presence of 50% RRL, 30 mM HEPES–KOH (pH 7.5), 50 mM KOAc, 1.0 mM Mg(OAc)_2_, 0.2 mM ATP, 0.2 mM GTP, 0.04 mM of 20 amino acids (Promega), 5 mM DTT, and 1% furimazine NanoLuciferase substrate (Promega; N113A). To initiate translation, 11 µL of reactions were preincubated at 30°C for 3 min before adding *in vitro* transcribed mRNA encoding Nanoluciferase (100 nM final concentration) to a final reaction volume of 12 µL. Translation kinetics were measured over time by recording NanoLuciferase luminescence using an Infinite m1000 pro microplate reader (Tecan) at 30°C for 15 min. The maximum translation rates (Max ΔRLU/Δsec) were determined as the peak values of the first derivative of the RLU curve, calculated using Prism 10 (GraphPad Software).

### Plunge freezing

For MCF-7 and RRL lysates, Quantifoil R2/1 holey-carbon grids coated with a thin layer of carbon (Electron Microscopy Service) were glow-discharged with 20 mA current with negative polarity for 30 s in a PELCO easiGlow glow discharge unit. The Vitrobot Mark IV (Thermo Fisher Scientific) was pre-equilibrated to 4 °C and 100% relative humidity and the blot force was set to zero. For all samples, 2.5 µl of lysate for each blotting was applied to the grid (with no waiting time), blotted 2 or 3 times for 3-7 seconds, force zero, and plunged into liquid-nitrogen-cooled liquid ethane. Grids were stored in liquid nitrogen.

For BSC-1 cells, Quantifoil gold (Au) mesh grids with a holey SiO_2_ film (R 2/2) were glow-discharged using an EMITECH K100X system, applying a negative coating current of 25 mA for 45 seconds. Following treatment, the grids were blotted from the reverse side and rapidly plunged into liquid ethane at -184°C using a Leica EM GP plunger (Leica Microsystems) maintained at 15°C with 85% relative humidity. Blotting times were set to 8 seconds. The vitrified grids were subsequently stored in liquid nitrogen within sealed boxes until further processing.

### Cryo-EM data collection and analysis

All data were collected at the UMass Chan Medical School cryo-EM facility. Data for all samples except for starved MCF-7 cell lysates were acquired using a Krios electron microscope (Thermo Fisher Scientific) operating at 300 kV, equipped with a Gatan Image Filter (slit width, 20 eV) and a K3 direct electron detector (Gatan). Data from starved MCF-7 lysates were collected on the Talos, 200 kV. Data collection was automated with SerialEM [116] using beam-image shift to collect multiple videos (for example, five videos per hole at four holes) at each stage position, and targeting 0.7 μm to 2 μm underfocus.

The starved MCF-7 cells dataset contained ∼10,000 movies collected with a total exposure of 40 e^−^/Å^−2^ and yielded 88,560 particles. For MCF-7 cells with no stress, the dataset comprised ∼10,000 movies with an exposure of 46 e^−^/Å^−2^ yielded 89,484 particles. The movie frames were aligned during data collection using IMOD [117] to decompress frames, apply the gain reference, and correct for image drift and particle damage. 2DTM was performed on parallelized Nvidia GPU processors (RTX A5000 and A6000), with the match_template program [21] implemented in *cis*TEM on motion-corrected images (pixel size 0.83 Å for non-starved and 0.87 Å for starved cells), using an in-plane angular step of 2.5° and an out-of-plane step of 3.5°, with the defocus search turned on for micrographs with thicker ice. To generate the template, eIF6 was removed from PDB 7OW7, and the remaining 60S subunit was converted to a density map using EMAN2 [112] with the box size matching the stack, and the B-factor of 80 Å² was applied using bfactor.exe from Frealign. Template-matched x,y coordinates, Euler angles, and defocus values from CTFFIND5 [118] were extracted using the MT package in the *cisTEM* GUI, generating a .star file. The MT package was used to import the .star file, and the *cis*TEM refine package was utilized to prepare the particle stack with a box size of 608 x 608 x 608 pixels. The refine package stacks were exported into the Frealign format for maximum-likelihood classifications (**Figure 2—figure supplement 3, 4**). Initial 3D reconstructions were generated using the Generate 3D tool in *cis*TEM. Further refinement of particle alignments included one cycle of x,y shifts, followed by a cycle of Euler angles and another cycle of defocus and beam tilt refinement, all performed manually using the *cis*TEM GUI.

Four datasets were collected from RRL samples: two datasets from one grid prepared with mRNA, and two datasets from one grid without mRNA. The first two datasets comprised 13,127 and 14,697 movies (26 frames, with an exposure of 1.515 e^-^ Å^-^² per frame, resulting in a total exposure of 39.39 e^-^ Å^-^² per sample or exposure of 1.538 e^-^ Å^-^² per frame, resulting in a total exposure of 39.35 e^-^ Å^-^² per sample, respectively). The data were combined to comprise 27,924 movies to be processed with 2DTM. The datasets for RRL samples without mRNA contained 29,855 movies (dataset A: 26 frames, with an exposure of 1.5167 e^-^ Å^-^² per frame, resulting in a total exposure of 39.4338 e^-^ Å^-^² per sample) and 44,983 movies (dataset B: 22 frames, with an exposure of 1.3745 e^-^ Å^-^² per frame, resulting in a total exposure of 30.238 e^-^ Å^-^² per sample). All data collections targeted 0.7 μm to 2 μm underfocus. 2DTM was performed with the match_template program [21] implemented in *cis*TEM on 2× binned images (pixel size 1.66 Å). To generate the template, eIF6 was removed from PDB 7OW7, and the remaining 60S subunit was converted to a density map using EMAN2 [112] with the box size matching the stack, and the B-factor of 200 Å² was applied using bfactor.exe from Frealign. The dataset was divided into blocks of 10,000 images (for each data collection) for 2DTM searches, yielding 209,874 particles from 27,924 movies of RRL with mRNA, 356,530 (dataset A) and 512,697 (dataset B) identified targets using an in-plane angular step of 3.5° and an out-of-plane step of 4°, without defocus search. Template-matched x,y coordinates, Euler angles, and defocus values from CTFFIND5 [118] were extracted using the MT package in the *cis*TEM GUI, generating a .star file. These .star files were subsequently merged with an in-house Python script, and the 2× binned image paths were replaced with the corresponding 1× binned image paths. The MT package was used to import the updated .star file, and the *cis*TEM refine package was utilized to prepare the particle stack with a box size of 800 x 800 x 800 pixels. The refine package stack was exported into a Frealign v9.11 [109] format for 3D classification (**Figure 3—figure supplement 2**).

Initial 3D reconstructions for the parent maps, were generated using the Generate 3D tool in *cis*TEM. Further refinement of particle alignments included one cycle of x,y, followed by a cycle of Euler angles and another cycle of defocus and beam tilt refinement, all performed manually using the *cis*TEM GUI.

Data from BSC-1 samples were acquired using a Krios electron microscope (Thermo Fisher Scientific) operating at 300 kV, equipped with a Gatan Image Filter (slit width, 20 eV) and a K3 direct electron detector (Gatan), with a target defocus range of 1.0–1.5 μm. Automated data collection was conducted using SerialEM [116], employing beam-image shift to acquire multiple movies (5 per hole across 9 holes) at each stage position. Zero-loss peak (ZLP) refinement was performed every 90 minutes at a unique location to avoid dark areas. The dataset from BSC-1 cells consisted of 24,550 movies, each containing 30 frames, with an exposure of 1.02 e^-^ Å^-^² per frame, resulting in a total exposure of 30.5 e^-^ Å^-^² per sample.

MotionCor2 [119] was used for drift correction, gain reference application, exposure weighting, and binning of super-resolution pixels by factors of 2 and 4 to yield final pixel sizes of 0.83 Å and 1.66 Å, respectively. Contrast transfer function (CTF) estimation was performed using CTFFIND5 [118] through the *cis*TEM graphical user interface (GUI) [38]. For 2D template matching (2DTM), 3D templates were generated from a trimmed 5LZV PDB model containing only the 60S ribosomal subunit, using the simulate program [120] from *cis*TEM to produce a 3D volume. A pixel size of 1.66 Å and a linear scaling factor of PDB B-factor of 1 were applied.

2DTM was performed with the match_template program [21] implemented in *cis*TEM on 4× binned images (pixel size 1.66 Å). The dataset was divided into blocks of 2,000 images for 2DTM searches, yielding 525,827 identified targets using an in-plane angular step of 1.5° and an out-of-plane step of 2.5°, without defocus search. Template-matched x,y coordinates, Euler angles, and defocus values from CTFFIND5 were extracted using the MT package in the *cis*TEM GUI, generating a .star file. These .star files were merged using an in-house Python script, and the 4× binned image paths were replaced with the corresponding 2× binned image paths. The MT package was used to import the updated .star file, and the *cis*TEM refine package was utilized to prepare the particle stack with a box size of 560 x 560 x 560 pixels.

Initial 3D reconstructions were generated using the Generate 3D tool in *cis*TEM. Further refinement of particle alignments included one cycle of x,y, followed by a cycle of Euler angles and another cycle of defocus and beam tilt refinement, all performed manually using the *cis*TEM GUI. The final alignment parameters and defocus values were exported in Frealign format.

### Cryo-EM data classification

#### MCF-7 lysates

Particle classifications were performed in Frealign v9.11 [109]. For both MCF-7 datasets, box size was 608 x 608 x 608 pixels. To speed up the processing, binned image stacks (e.g. 8×, 4× or 2×) were prepared using resample.exe part of the Frealign v9.11 distribution. Focused 3D maximum-likelihood classification into 24 classes (using the high-resolution limit of 16 Å and the 8× binned stack, 100-150 rounds) with an 80-Å focus mask covering the A site and the GTPase-activating center of the ribosome (x, y, z = 261.73, 312.71, 208.96) resolved ribosomes with different occupancies of translation factors (**Figure 2—figure supplement 3, 4**).

Classes were merged using merge_classes.exe from Frealign v9.11, applying a class occupancy threshold of 0.50 and a score of 0, followed by un-binning of particles using Frealign v9.11. Additional subclassifications were performed but did not result in additional high-resolution classes beyond those appearing in the initial classification.

#### RRL

The box size for RRL datasets was set to 800 x 800 x 800 pixels. To speed up processing, binned image stacks (e.g. 8×, 4× or 2×) were prepared using resample.exe, part of the Frealign v9.11 distribution. Focused 3D maximum-likelihood classification into 40 classes (using the high-resolution limit of 16 Å and the 8× binned stack, 100 rounds) with an 80-Å focus mask covering the A site and the GTPase-activating center of the ribosome (x, y, z = 309.98, 345.63, 232.03) resolved different ribosome states (**Figure 3—figure supplement 2**).

For RRL with mRNA, initial classification of 209,874 particles yielded two low-resolution (junk) classes with 13,791 particles (6.6%) that were removed from further classification. Out of 9 classes corresponding to 60S subunits, 2 classes contain eEF2 and the remaining ones represent 60S without additional factors (Table S5). 31 classes were 80S ribosomes: the predominant 19 classes (99,843 particles) contain eIF5A, SERBP1 and eEF2, and 3 classes (15,065 particles) represent the codon sampling states with eEF1A and A/T tRNA. One class with 4036 particles has tRNA in the A and P sites and additional density in the A-site, which improved upon merging this class with particles from RRL without mRNA that featured nearly identical density. The additional A-site density resembles DRG1 or DRG2 (**Figure 3—figure supplement 3B**). Six classes (22,093 particles) featured eEF2, although different density levels at factor-binding sites suggested that the cryo-EM maps are insufficiently separated. These 6 classes were merged (with criteria >50% occupancy and Score >0), resulting in a 17,529-particle stack and then subclassified to 8 different classes with a mask on the P-site. This classification resolved 3 predominant complexes: with eEF2, IFRD2 and LARP1 in the mRNA tunnel, with eEF2 and two tRNAs and with eEF2, E-tRNA with SERBP1. For RRL without mRNA, classification of dataset A was performed with the same mask as in the above classification.

To achieve higher resolution, classes representing the same functional states in different RRL datasets were merged using merge_classes.exe from Frealign, applying a class occupancy threshold of 0.50 and the Score of 0, followed by 3D 1x binned reconstruction of particles using FrealignX or *cis*TEM. Initial 3D reconstructions were generated using the Generate 3D tool in *cis*TEM. Further refinement of particle alignments including one cycle of x,y, followed by a cycle of Euler angles and another cycle of defocus and beam tilt refinement, were performed using the *cis*TEM GUI.

For 80S with IFRD2 and LARP1, all representative classes from different datasets were merged in IMOD (71,194 particles). Using a 3D mask for eEF2 (eEF2 from PDB 6MTE was aligned to a 8×-binned map containing eEF2 and then a 3D map was created based on the eEF2 model using molmap in ChimeraX [121], then masked using vop onesmask #newmap onGrid #oldmap and saved as an mrc file), this stack (binned to match the 8×-binned stack) was classified into two classes, resulting in two maps, both containing IFRD2 and LARP1, and either with or without eEF2. For 80S with eEF2 and CCDC124, particles were extracted with >50% occupancy and score >0, and merged (27,910 particles). This stack was 4× binned and subclassified into 6 classes using a 45 Å mask on the P and E sites. For 60S with eEF2, classes with eEF2 from different datasets were merged in IMOD (38,579 particles) and classified using a 34 Å focus mask on eEF2 domain IV, resulting in 60S with open or compact eEF2.

#### BSC-1

An initial 3D maximum-likelihood classification (without particle alignment) was performed in Frealign v9.11 on the 4× binned stack, using 100 classes and 14 classification cycles [109]. The high-resolution limit for classification was set to 14 Å. Classes corresponding to equivalent states were then merged using an in-house Python script that selects particles with occupancy greater than 50%. This script preserves the file paths and generates a particle alignment file (.star) compatible with cisTEM [21]. The latter was used to generate the particle stack (.mrc) in cisTEM, which also allowed for refinement of particle alignments for each class in cisTEM. The script is available at https://github.com/GrigorieffLab/yafw. Subclassification was performed using a focus mask and 4× binned data (**Figure 2—figure supplement 1**), with Frealign v9.11. 3D reconstructions with the unbinned data (physical pixel size of 1.66 Å) were generated using either cisTEM or FrealignX.

### Model building and refinement

Cryo-EM structure of (GR)_20_-bound *O. cuniculus* 80S ribosome (PDB: 7TOR [111]), omitting (GR)_20_ and P-tRNA, was used as the starting model for structure refinement into all maps (Table S4). 60S, 40S head, 40S body were rigid-body fitted into each cryo-EM map using ChimeraX [121] based on PDB 6MTD as a reference structure for head swiveled models. For eEF2 (P13639.EF-HUMAN), IFRD2 (Q12894-IFRD2-HUMAN), uL6 (P32969-RL9-HUMAN), ul14 (P62829.RL2 3_HUMAN), eL30 (P62890.RL30_RAT), uS19 (P62841.RS15_HUMAN), and eS28 (P62857.RS28-HUMAN), AlphaFold [35] models were fitted into density, using ChimeraX [121]. The EBP1-ES27L structure was modelled from PDB: 7BHP [52], LARP1 from PDB: 8XP2 [32], CCDC124 from PDB: 6Z6L [34], E-tRNA from PDB: 7OSM [87], uL1 and eS12 from PDB: 6MTD. To model open and closed conformations of eEF2 in 60S complexes (Fig. 3K, L), cryo-EM maps from both conformations were softened by applying the B-factor of 100 Å^2^ and rigid-body fitted in Chimerax [121]. After rigid-body fitting and manual modeling, structural models were refined using phenix.real_space_refine [122–124], yielding structures with good stereochemistry and fits into corresponding cryo-EM maps (Table S4).

## Data availability

Structural models generated in this study have been deposited in the RCSB Protein Data Bank under the following accession codes: **11IQ** (Rabbit 80S ribosome with eEF2, eIF5a and SERBP1), **11HG** (Rabbit 80S ribosome with eEF2, CCDC124 and LARP1), **11JJ** (Rabbit 80S ribosome with eEF2, IFRD2 and LARP1), **11KH** (Rabbit 80S ribosome with IFRD2 and LARP1), **11HV** (Rabbit 60S with eEF2, domain IV open), **11HE** (Rabbit 60S with eEF2, domain IV closed).

Cryo-EM maps used to generate these models have been deposited in the Electron Microscopy Data Bank under the following accession codes: **EMD-75724** (Rabbit 80S ribosome with eEF2, eIF5a and SERBP1), **EMD-75689** (Rabbit 80S ribosome with eEF2, CCDC124 and LARP1), **EMD-75740** (Rabbit 80S ribosome with eEF2, IFRD2 and LARP1), **11KH** (Rabbit 80S ribosome with IFRD2 and LARP1), **EMD-75704** (Rabbit 60S with eEF2, domain IV open), **EMD-75687** (Rabbit 60S with eEF2, domain IV closed).

## Acknowledgments

We thank Chen Xu, Kangkang Song and Christna Ouch for the help with data collection at the cryo-EM facility at UMass Chan Medical School; Johannes Elferich for assistance with 2DTM data processing; members of the Grigorieff and Korostelev laboratories for helpful discussions and comments on the manuscript. This study was supported by the Howard Hughes Medical Institute (HHMI) to N. G., the US National Institutes of Health grants CHEETAH U54AI170856 and R35GM127094 to A.A.K.

## Author contributions

Z.S. prepared cell cultures and cryo-EM samples, collected and analyzed cryo-EM data, prepared illustrations, and drafted the manuscript; X.Z. prepared cell cultures and cryo-EM samples, collected and analyzed cryo-EM data, and prepared illustrations; C-Y. H. performed biochemical experiments, analyzed and interpreted cryo-EM data, and prepared illustrations; A.B.L, S.D. and E.S. analyzed and interpreted cryo-EM data; N.G. supervised data analyses and secured funding; A.A.K. conceptualized the study, supervised data analyses, drafted the manuscript and secured funding. All authors contributed to data interpretation and manuscript finalization.

## Notes

### Competing Interest Statement

The authors have declared no competing interest.

### Summary of Updates

In this revised version, we have addressed the reviewers comments. We also added the method workflow and several additional supplementary figures. The PDB IDs are now available as well.

