## Supplementary Information for "*In extracto* cryo-EM reveals eEF2 as a major hibernation factor on 60S and 80S particles"

**Contents:**

Supplementary Table 1-5
Supplementary Figures 1-9
Supplementary References

**Table S1.** Particle picking strategies for the RRL+angiogenin dataset.

| Strategy | Particles picked | Best resolution of classes | Classes observed |
|---|---|---|---|
| *cis*TEM (Best 600 micrographs, CTF fit 2.5-3.2 Å) | ~8000 | >30 Å, would not align | none |
| *cis*TEM (full stack)/ Frealign | 272,478 | >30 Å, would not align | none |
| Relion v3 (full stack, picked with crYOLO trained on best 6216 particles from handpicking) | 72,431 or subsets | >30 Å, would not align | none |
| Relion v3 (full stack, LoG picked and culled for bad ice and by lnCtfMaxRes ={1,4}) | 83,055 | 14 Å | 1) 80S+Ang+TC |
| Relion v3 (full stack, LoG picked and culled for bad ice and CTF, choosing high-resolution 2D classes) | 52,580 | < 6.6 Å (binned) | 1) 80S+Ang+TC |
| Relion v3 (full stack, LoG picked and culled of bad ice, bad CTF, bad 2D classes, and selecting highest resolution 3D class for 2$^{nd}$ round of subclassification) | 16,848 | 3.2 Å | 1) 80S+Ang+TC<br>2) 80S+eEF2 stalled<br>3) 60S |
| Hand picking (***best*** 600 micrographs, refine picks, recenter...)/Frealign v9 | 6216 | 3.5 Å | 1) 80S+Ang+TC<br>2) 80S+Ang+weakTC<br>3) 80S+eEF2 stalled<br>4) 60S |
| 2DTM (~1600 micrographs)/ Frealign v9 | 37,859 | 3.1 Å | 1) 80S+Ang+TC<br>2) 80S+Ang<br>3) 80S+TC<br>4) 80S+2tRNAs+eEF2<br>5) 80S+eEF2 stalled<br>6) 60S |
| 2DTM (full stack)/ Frealign v9 | 88,488 | 3.0 Å | 1) 80S+Ang+TC with substates<br>2) 80S+Ang<br>3) 80S+TC with substates<br>4) 80S+2tRNAs+eEF2 with substates<br>5) 80S+eEF2 stalled<br>6) 60S |

**Table S2.** Ribosome states observed in lysates from different cell types.

| **Ribosome status**<br><br>*Cell type* | **Elongating (with mRNA and tRNAs and/or eEF1A/eEF2)** | **Hibernating** | **60S** | **Other classes** | **# particles and Resolution (average initial reconstruction)** |
|---|---|---|---|---|---|
| *MCF-7 lysate* | 71.9% | 18% | 9.9% | - | 89,484-2.7 Å |
| *MCF-7 starved, lysate* | 57.6% | 27.3% | 14.9% | - | 88,560-2.6 Å |
| *BSC-1 lysate* | 58.3% | 32.6% | 6.5% | 2.2 % (ABCE1/eRF1) | 525,827-2.2 Å |
| *RRL with mRNA* | 12.4% | 58.6% | 26% | 3% (initiation) | 209,874-2.2 Å |
| *HeLa cells (2DTM/cryo-EM; Zheng et al, 2024)[1]* | 70.1% | 29.9% | Unknown | - | 2.2 Å |
| *HEK293 cells, ER-bound 80S (cryo-ET, Gemmer et al, 2023)[2]* | 90% | 9.9% | Unknown | - | 4.4 Å |

**Supplementary Table 3.** Rotation of the 40S small ribosomal subunit head or body relative to nonrotated 80S ribosomes: with an accommodation-like eEF1A•aa-tRNA (PDB ID ; 5LZS [3]) or with tRNAs in three classical sites (this work).

| | Relative to non-rotated 80S with eEF1A•aa-tRNA and P-tRNA (PDB 5LZS) | | Relative to non-rotated 80S with 3 tRNAs: A, P, E (this work) | |
|---|---|---|---|---|
| | **Head** | **Body** | **Head** | **Body** |
| **eEF2+eIF5A+SERBP1** | 19.7° | 5.2° | 16.0° | 4.0° |
| **eEF2+CCDC124+LARP1** | 17.8° | 0.8° | 14.2° | 4.8° |
| **eEF2+IFRD2+LARP1** | 17.8° | 0.9° | 14.2° | 4.8° |
| **IFRD2+LARP1** | 18.0° | 0.2° | 14.5° | 4.3° |

**Supplementary Table 4.** Cryo-EM data collection and model refinement statistics for 80S and 60S complexes from RRL.

| | 80S•eEF2 •eIF5a•SERBP1 | 80S•eEF2 •IFRD2•LARP1 | 80S•IFRD2 •LARP1 | 80S•eEF2 •CCDC124•LARP1 | 60S•eEF2 (open) | 60S•eEF2- (closed) |
|---|---|---|---|---|---|---|
| **RCSB PDB ID** | 11IQ | 11JJ | 11KH | 11HG | 11HV | 11HE |
| **EMDB Accession ID** | EMD-75724 | EMD-75740 | EMD-75768 | EMD-75689 | EMD-75704 | EMD-75687 |
| **Data collection and processing** | | | | | | |
| Nominal magnification | 60,241x | 60,241x | 60,241x | 60,241x | 60,241x | 60,241x |
| Voltage (kV) | 300 | 300 | 300 | 300 | 300 | 300 |
| Electron exposure ($e^{-}/Å^2$) | 30 | 39 | 39 | 30 | 39 | 39 |
| Defocus range (μm) | 0.7-2 | 0.7-2 | 0.7-2 | 0.7-2 | 0.7-2 | 0.7-2 |
| Pixel size (Å) | 0.83 | 0.83 | 0.83 | 0.83 | 0.83 | 0.83 |
| Symmetry imposed | C1 | C1 | C1 | C1 | C1 | C1 |
| Initial particle images (no.) | 512,697 | 1,079,101 | 1,079,101 | 512,697 | 1,079,101 | 1,079,101 |
| Final particle images (no.) | 198,920 | 16,727 | 10,932 | 8,501 | 15,960 | 22,323 |
| Map resolution (Å)** | 2.4 | 3.0 | 3.2 | 2.7 | 2.9 | 2.9 |
| FSC threshold | 0.143 | 0.143 | 0.143 | 0.143 | 0.143 | 0.143 |
| **Refinement** | | | | | | |
| Initial model used (PDB code) | 7TOR | 7TOR | 7TOR | 7TOR | 7TOR | 7TOR |
| **Model resolution (Å)*** | | | | | | |
| FSC threshold | 0.143 | 0.143 | 0.143 | 0.143 | 0.143 | 0.143 |
| Correlation Coefficient (cc_mask)* | 0.84 | 0.80 | 0.79 | 0.79 | 0.76 | 0.75 |
| **Model composition*** | | | | | | |
| Non-hydrogen atoms | 224497 | 224042 | 218411 | 221974 | 147591 | 147624 |
| Protein residues | 13401 | 13271 | 12433 | 12998 | 8155 | 8159 |

| | | | | | | |
|---|---|---|---|---|---|---|
| RNA residues | 5467 | 5497 | 5539 | 5497 | 3825 | 3825 |
| **R.m.s. deviations from ideal*,§** | | | | | | |
| Bond lengths (Å) | 0.002 | 0.003 | 0.003 | 0.002 | 0.009 | 0.004 |
| Bond angles (°) | 0.62 | 0.61 | 0.76 | 0.60 | 0.88 | 0.64 |
| **Validation*** | | | | | | |
| MolProbity score | 1.87 | 1.92 | 1.96 | 1.95 | 2.45 | 1.98 |
| Clashscore | 6.12 | 7.02 | 7.30 | 7.70 | 8.77 | 6.60 |
| **Ramachandran plot*** | | | | | | |
| Favored (%) | 97.68 | 97.71 | 97.36 | 97.74 | 95.94 | 96.88 |
| **Validation (RNA)*** | | | | | | |
| Good sugar pucker (%) | 98.82 | 98.83 | 98.91 | 98.83 | 98.69 | 98.77 |
| Good backbone (%)# | 99.24 | 99.23 | 99.01 | 99.23 | 99.08 | 99.32 |

*Statistics calculated in Phenix v.2.0[4].

**FSC_part from Frealign v. 9.11[5].

#RNA backbone suites that fall into recognized rotamer conformations defined by MolProbity[6].

**Supplementary Table 5.** Percentages of particles corresponding to 80S or 60S classes in RRL with NLuc mRNA and in RRL without NLuc mRNA.

| Ribosome state | Factors# | RRL+mRNA | RRL-mRNA |
|---|---|---|---|
| **Initiation** | eIF5B | 2.4% | 5.0% |
| **Elongation** | eEF1A+3 tRNA | 7.7% | 4.3% |
| | eEF2+2 tRNA | 1.0% | X* |
| | A/P and P/E tRNAs | X | 1.2% |
| | DRG1/2 | 2.0% | 0.7% |
| **Hibernation** | eEF2+eIF5A+SERBP1 | 51% | 46.5% |
| | eEF2+E-tRNA+SERBP1 | 5.7% | 2.0% |
| | eEF2+CCDC124+LARP1 | 1.1% | 5.4% |
| | eEF2+IFRD2+LARP1 | 1.0% | 3.1% |
| | IFRD2+LARP1 | X | 1.7% |
| **60S** | None | 19.2% | 26.4% |
| | eEF2 | 8.8% | 3.5% |

### EBP1 was identified in all the classes and is not specified in the table
*X denotes the states that were absent after 3D classifications

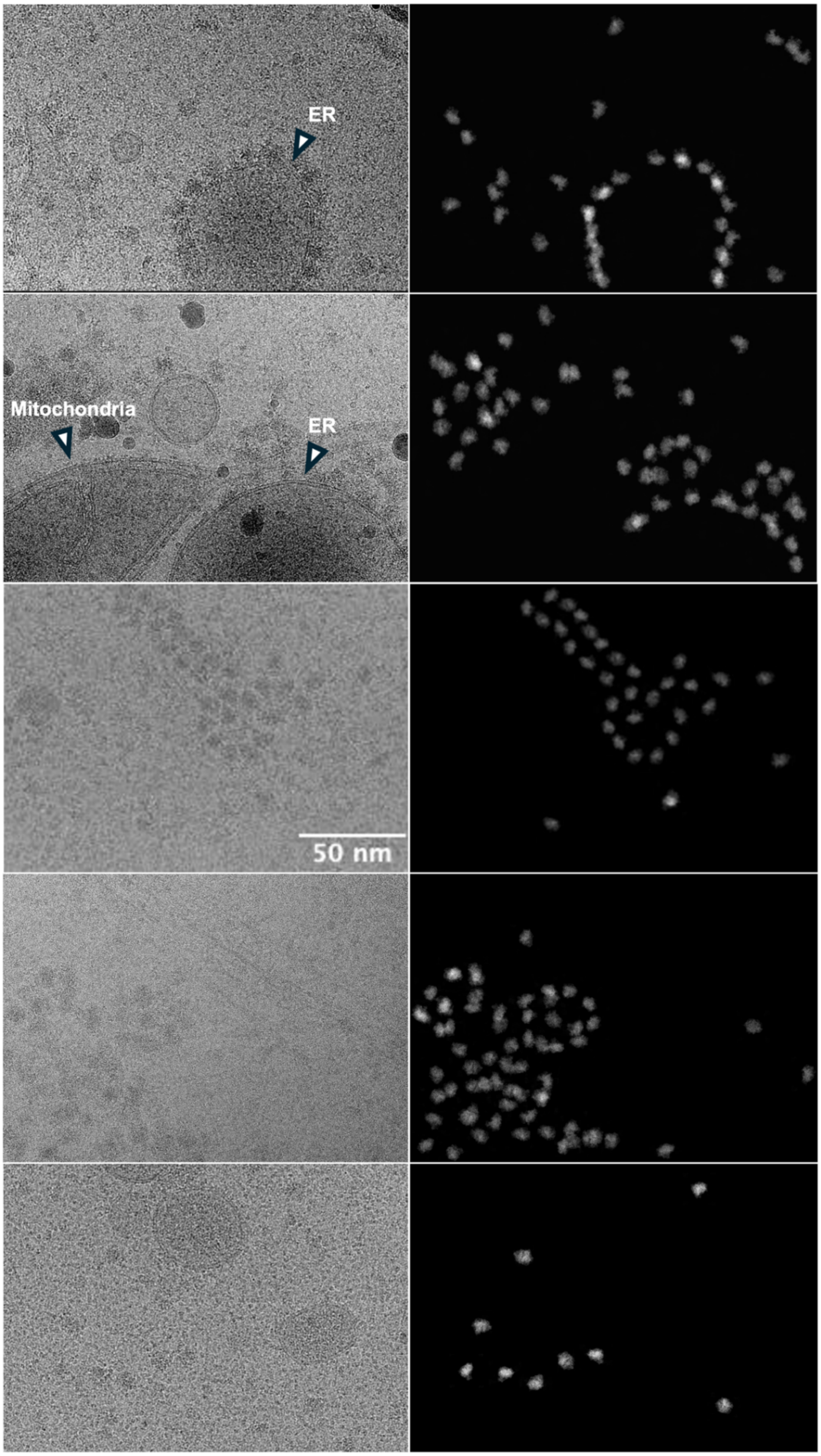


**Figure 1—figure supplement 1.** Examples of micrographs showing cellular context in BSC-1 lysates. Scale bar for all micrographs is 50 nm.

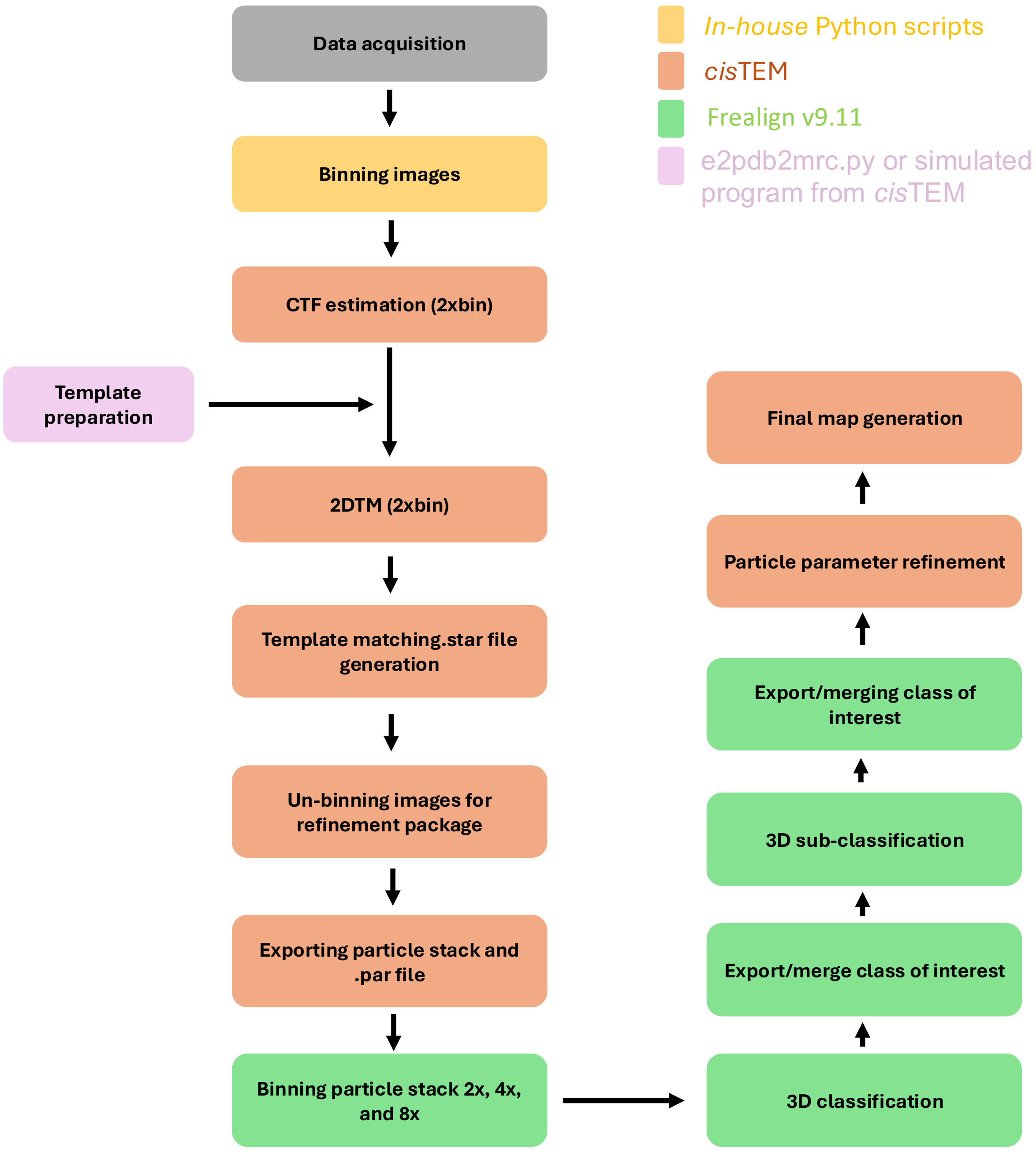


**Figure 1—figure supplement 2. Data processing workflow**. Micrographs were binned using in-house Python script and then contrast transfer function (CTF) parameters were estimated from 2× binned micrographs using *cis*TEM, followed by 2D template matching (2DTM) on binned data. To prepare the templates, we initially used both e2pdb2mrc.py and *cis*TEM's simulate program to generate templates (see Methods). Subsequently, we switched to exclusively using the simulate program since it preformed slightly better. Our goal in determining and optimizing search

parameters for 2DTM was a balance between particle search precision and computational cost (see methods). In practice, users must weigh the benefit of finer sampling against the substantial increase in runtime, particularly for large datasets. In some cases, the defocus search was turned off, and it was sufficient to use in-plane and out-of plane angular steps of approximately 4.5° and 3.5°, respectively, and binned images with ~2 Å/pixel, to still obtain enough matches to obtain high resolution reconstructions. To determine when defocus search was needed, micrographs were selected based on their fit resolution obtained during CTF estimation. From each dataset, we selected ten micrographs representing the highest and lowest fit resolutions. Template matching was then performed using identical parameters, once with defocus search enabled and once with it disabled. The number of detected particles for each micrograph under both conditions was compared. When a significant difference was observed, most commonly for icy micrographs with low fit resolution, we enabled defocus search for that group of images. We found these images/datasets appeared to have a higher background compared to *in-vitro* reconstituted samples but less than *in situ* samples The template matching results from these micrographs were subsequently combined with results from groups processed with defocus search disabled. After 2DTM, .star files were generated using *cis*TEM. The pixel size and directory to un-binned images was changed in this .star file, now called edited.star. A dummy project with a small group of micrographs was made and after performing 2DTM and generating a dummy.star file, the dummy.star file was substituted with edited.star. Then we generated the refinement package, corresponding particle stacks and .par files. Un-binning refers to generating particle stacks from the original micrographs at the original pixel size. Particle stacks and .par files were exported from *cis*TEM, and to speed up processing, binned image stacks (e.g. 2×, 4× or 8×) were prepared using resample.exe, which is part of the Frealign v9.11 distribution. Initial 3D classification was carried out in Frealign v9.11. For each resulting class, per-class particle stacks and .star files were generated and subsequent rounds of focused or global 3D classification were performed (see Methods). Software packages used at each step are indicated by color coding.

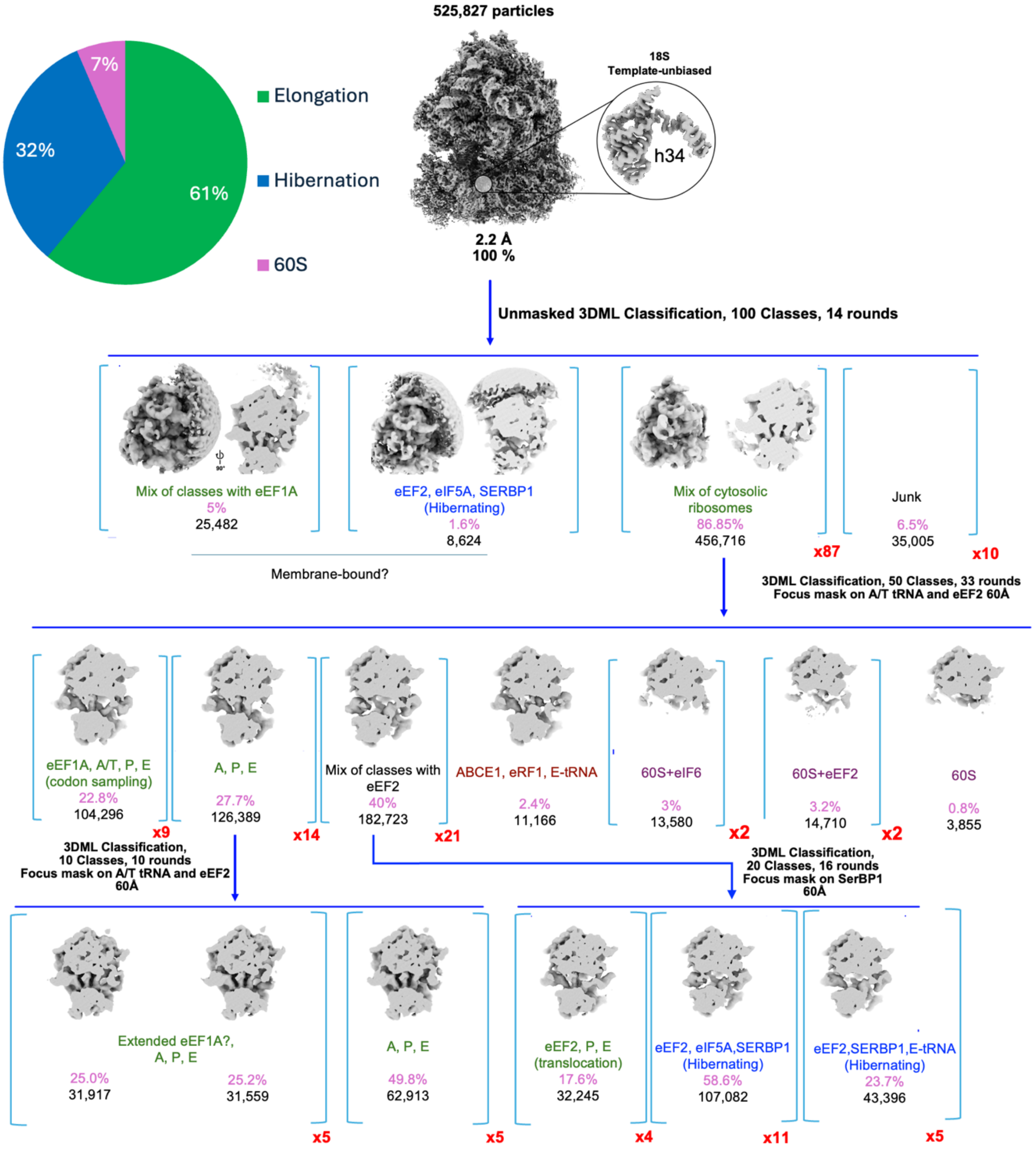


**Figure 2—figure supplement 1. Maximum likelihood 3D classification of the dataset collected from BSC-1 cell extracts.** Classification was performed on 8x binned stacks with particle alignment disabled.

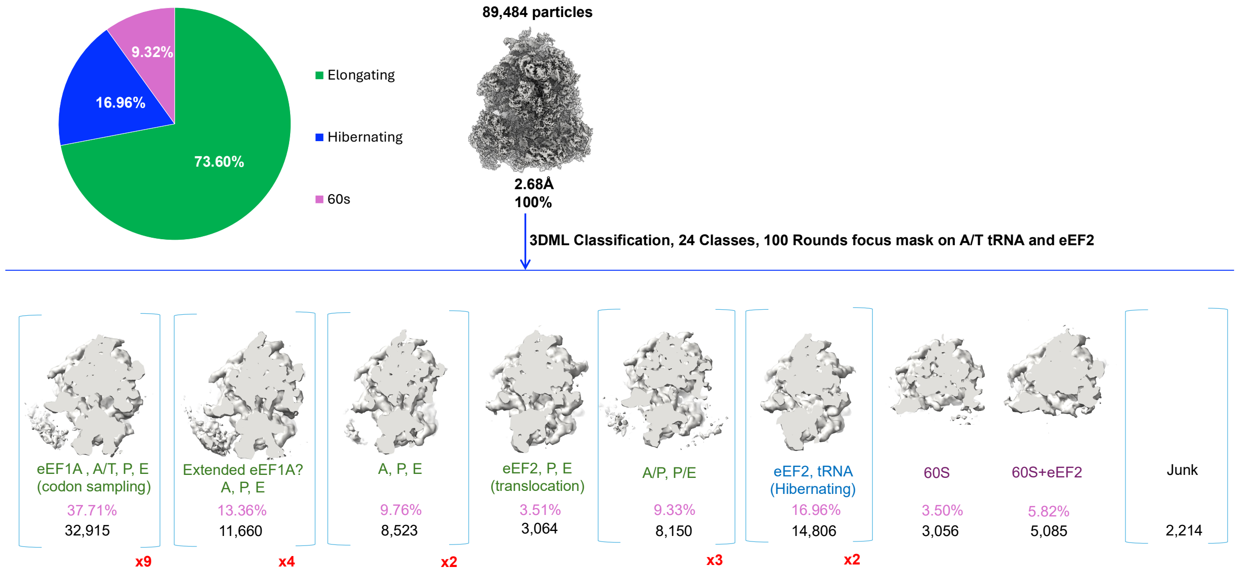


**Figure 2—figure supplement 2**. **Maximum-likelihood 3D classification of the nutrient-deprived MCF-7 extract dataset in Frealign v9.11 [5].**

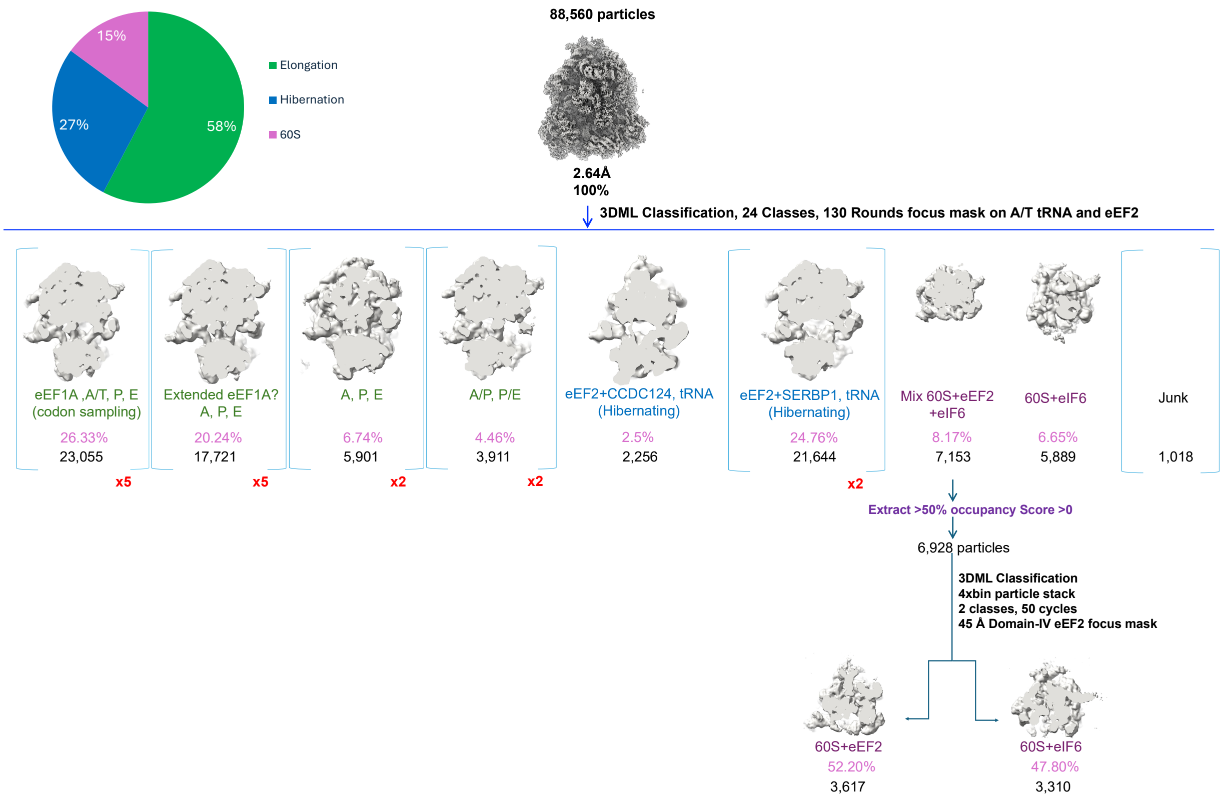


**Figure 2—figure supplement 3. Maximum-likelihood 3D classification of the MCF-7 extract dataset in Frealign v9.11.**

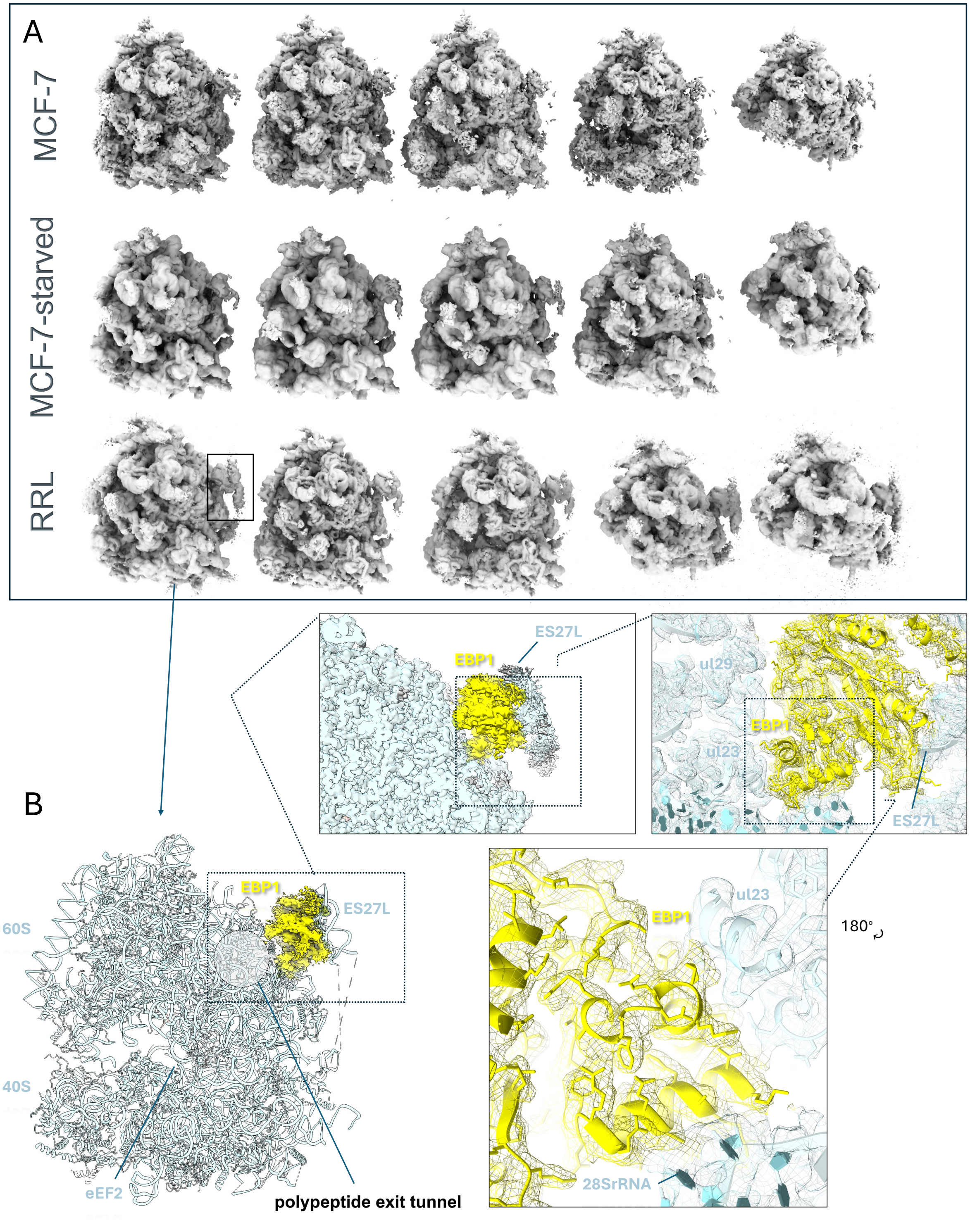


**Figure 3—figure supplement 1. Ribosome classes contain EBP1 bound next to the polypeptide exit tunnel**. **A**) Representative classes from different datasets featuring EBP1. **B**) A close-up view of EBP1 in RRL 80S ribosomes with eEF2, eIF5A, and SERBP1.

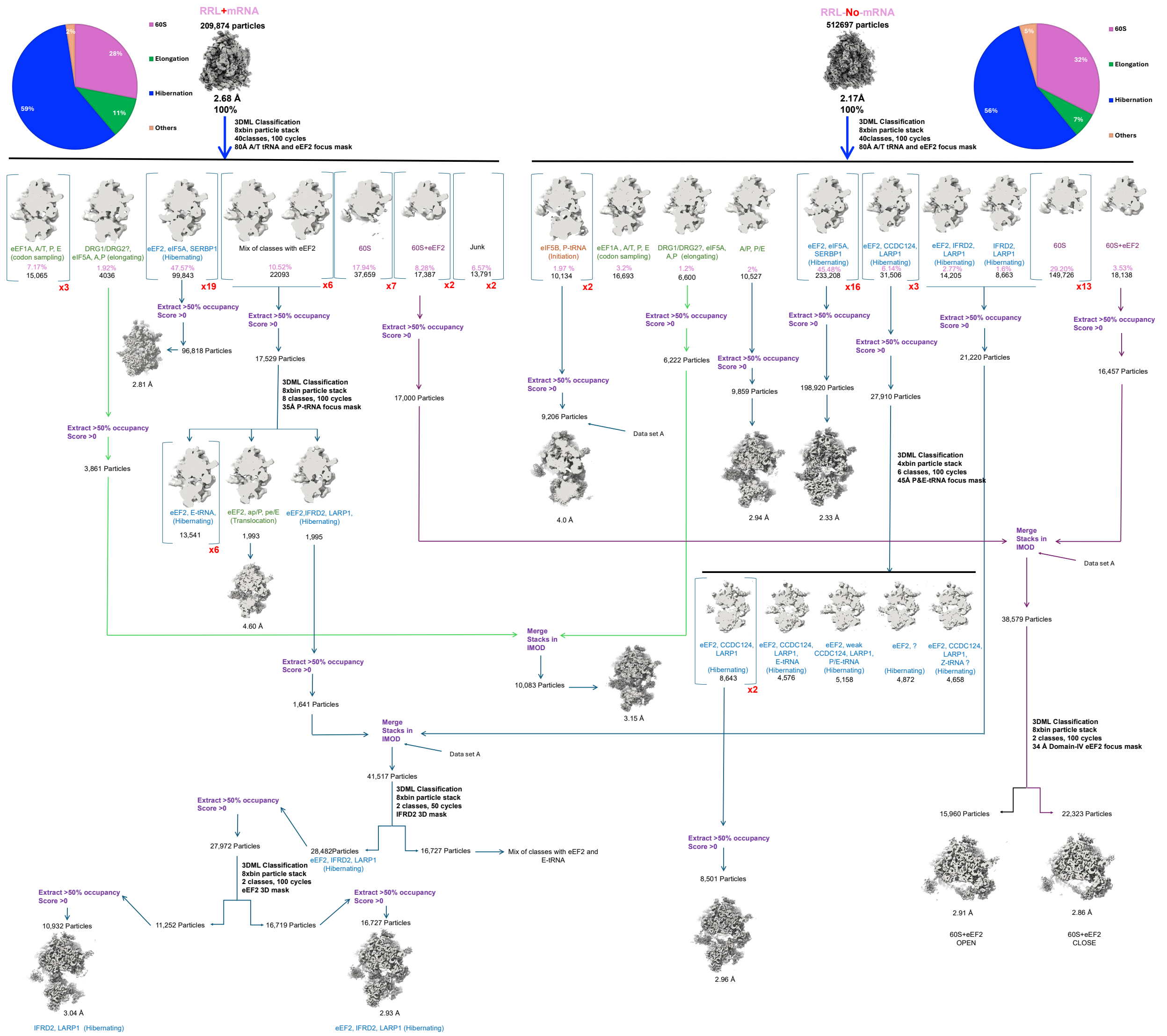


**Figure 3—figure supplement 2. Maximum-likelihood 3D classification of RRL datasets with or without nanoluciferase-encoding mRNA in Frealign v9.11.**

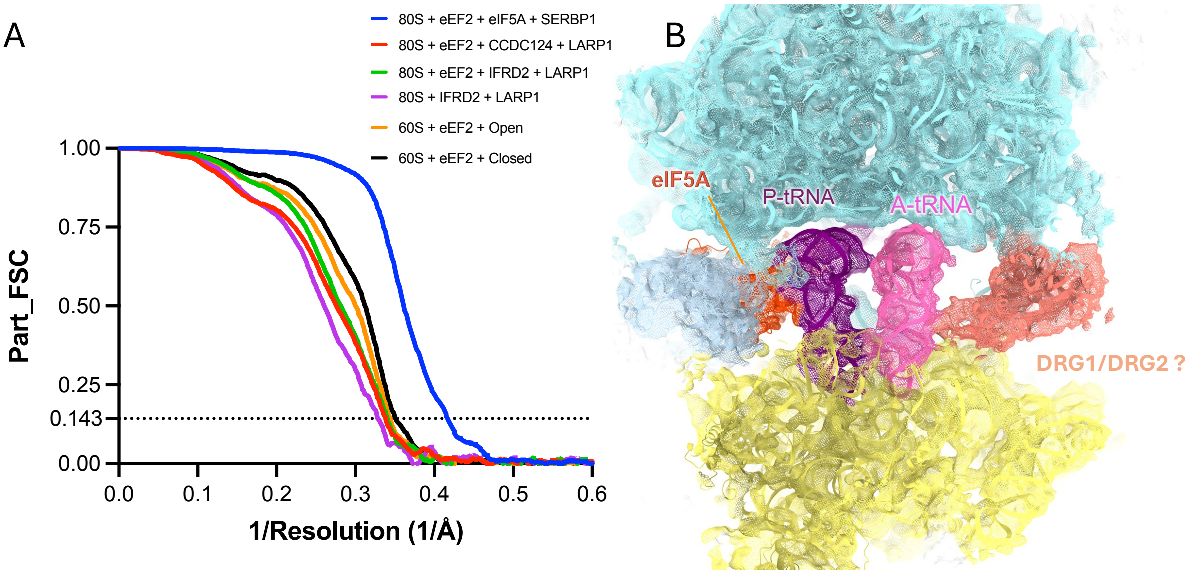


**Figure 3—figure supplement 3. A)** Fourier shell correlation (FSC) curves for RRL-derived maps (masked), as a function of inverse resolution. **B**) RRL 80S ribosomes with putative DRG1 (or DRG2), eIF5A in the E site, and tRNAs in the A and P sites.

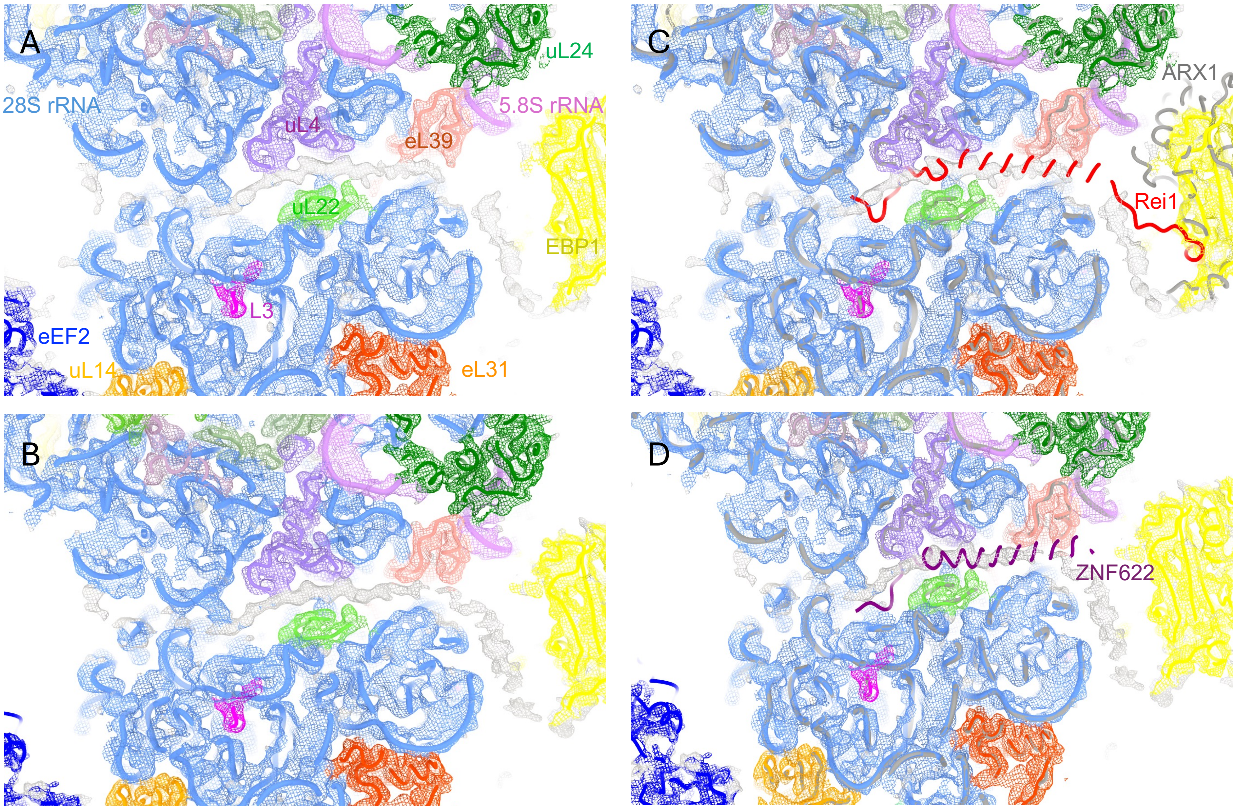


**Figure 3—figure supplement 4. 60S subunit polypeptide tunnel is occupied with an uncharacterized protein (RRL). A**) Cryo-EM map (mesh) and model of 60S with eEF2 open. Density corresponding to the tunnel protein is shown in grey. **B**) Cryo-EM map (mesh) and model of 60S with eEF2 closed. **C**) Superposition of the yeast 60S biogenesis factor Rei1 (PDB ID; 5APN [7]) with the tunnel density (grey). Rei1 is shown in red and the rest of the yeast 60S model is in gray. Cryo-EM maps were softened with the B-factor of 50 Å². 60S structure superpositions were performed in ChimeraX. **D**) Superposition of the mammalian 60S biogenesis factor ZNF622 (PDB ID; 9GMO [8]) with the un-identified tunnel density (grey). ZNF622 is shown in purple and the rest of the superposed 60S model is in gray.

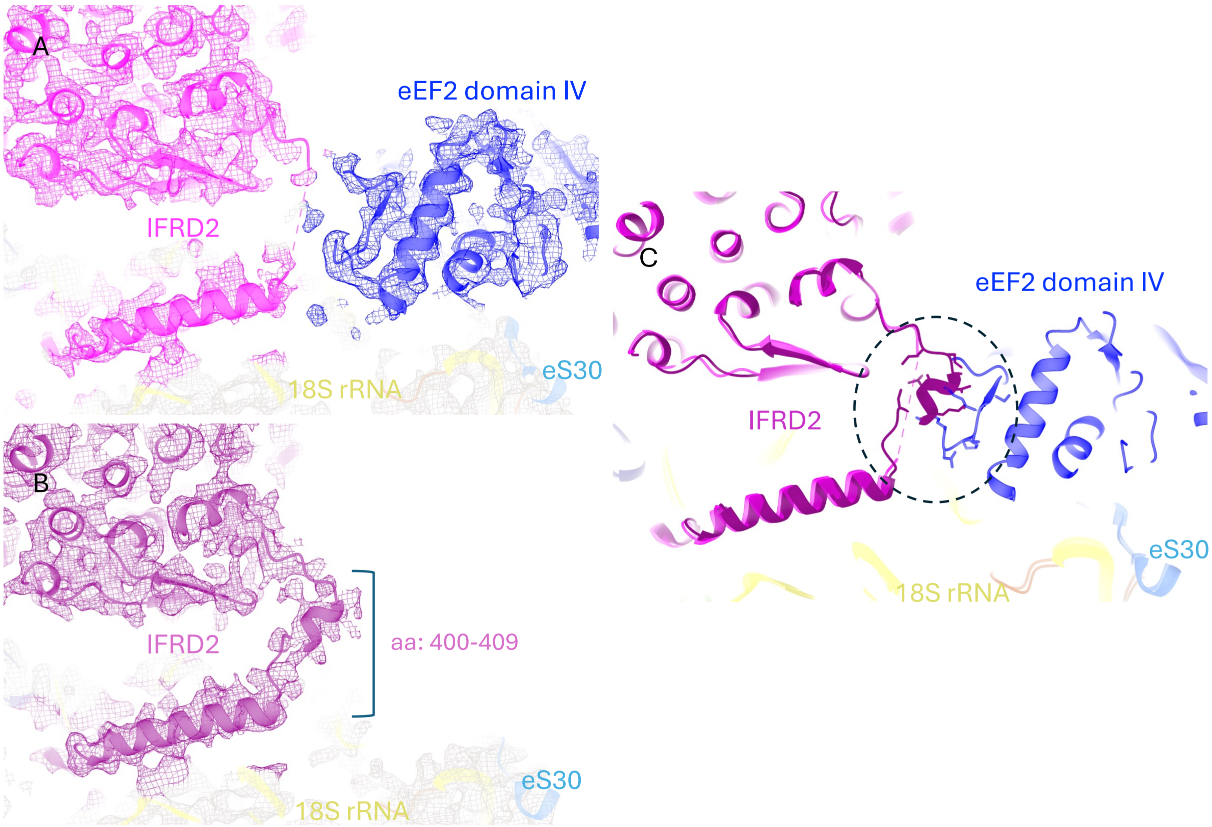


**Figure 3—figure supplement 5. The linker helix (aa: 400-409) of IFRD2 adopts different conformations in the presence and in the absence of eEF2**. **A, B**) Cryo-EM maps (mesh) and models of 80S with LARP1 and IFRD2, in the presence of eEF2 (A) and in the absence of eEF2 (B). **C**) Superposition of models from panel A and B. IFRD2 in the presence of eEF2 is shown in magenta and in the absence of eEF2 is shown in purple. The steric clash between the IFRD2 linker helix and domain IV of eEF2 is indicated by a circle.

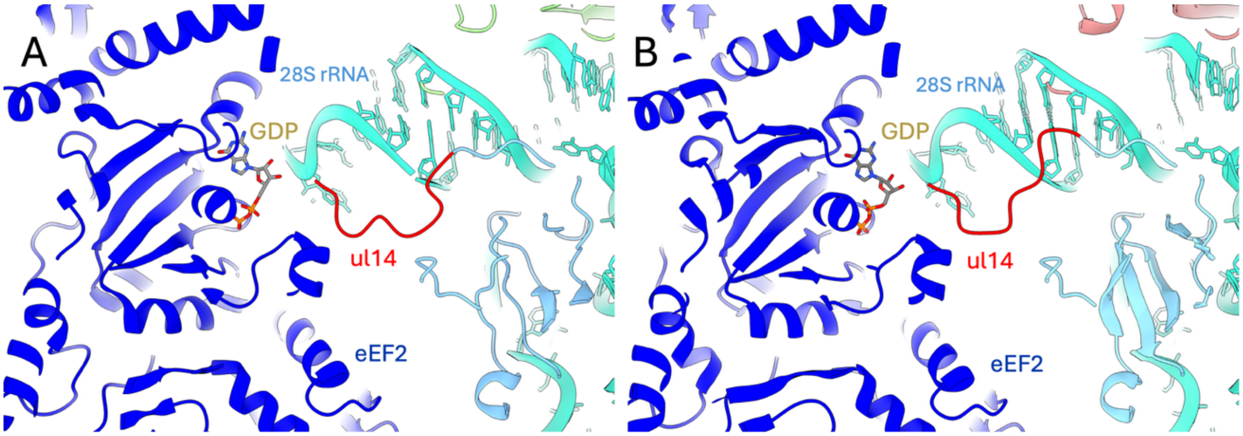


**Figure 1—figure supplement 1. N-terminal tail of uL14 in the presence of eEF2**. **A**) Structure with eIF5A, eEF2 and SERBP1 (this work, RRL). **B**) *G.gallus* eEF2 bound to translocation-like ribosomes (PDB ID; 8Q87 [9]).

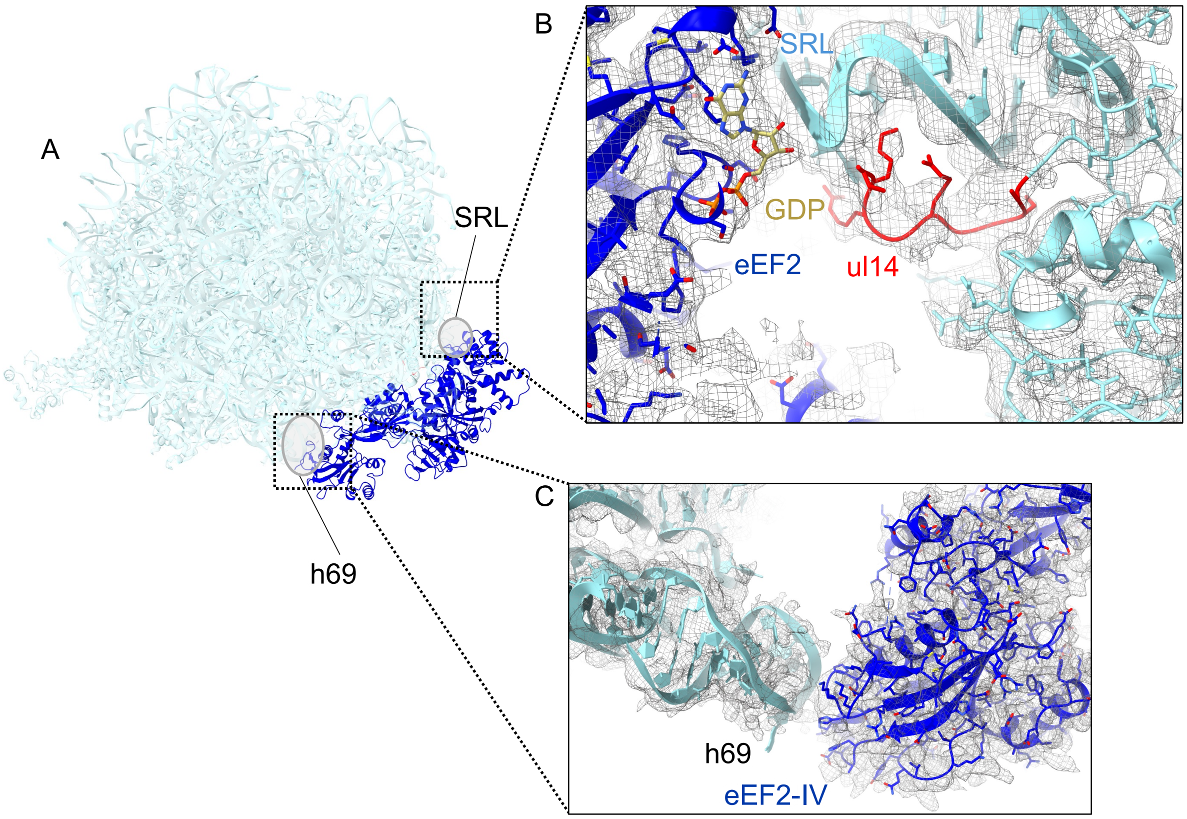


**Figure 4—figure supplement 2. Interaction of eEF2 with the isolated 60S subunit (RRL). A**) eEF2 in a compact conformation (relative to the second, more open conformation observed in this work). **B**) Interaction with the SRL and uL14. The cryo-EM map (mesh) was softened with a B-factor of 25 Å². **C**) Domain IV of eEF2 binds next to helix 69 of 28S rRNA. The map was softened with a B-factor of 30 Å².
